# Accelerating PERx Reaction Enables Covalent Nanobodies for Potent Neutralization of SARS-Cov-2 and Variants

**DOI:** 10.1101/2022.03.11.483867

**Authors:** Bingchen Yu, Shanshan Li, Takako Tabata, Nanxi Wang, G. Renuka Kumar, Jun Liu, Melanie M. Ott, Lei Wang

**Affiliations:** Department of Pharmaceutical Chemistry and the Cardiovascular Research Institute, University of California San Francisco, San Francisco, CA, USA; Gladstone Institutes, San Francisco, CA, USA; Department of Medicine, University of California San Francisco, San Francisco, CA, USA

## Abstract

The long-lasting COVID-19 pandemic and increasing SARS-CoV-2 variants demand effective drugs for prophylactics and treatment. Protein-based biologics offer high specificity yet their noncovalent interactions often lead to drug dissociation and incomplete inhibition. Here we developed covalent nanobodies capable of binding with SARS-CoV-2 spike protein irreversibly via proximity-enabled reactive therapeutic (PERx) mechanism. A novel latent bioreactive amino acid FFY was designed and genetically encoded into nanobodies to accelerate PERx reaction rate. After covalent engineering, nanobodies binding with the Spike in the down state, but not in the up state, were discovered to possess striking enhancement in inhibiting viral infection. In comparison with the noncovalent wildtype nanobody, the FFY-incorporated covalent nanobody neutralized both authentic SARS-CoV-2 and its Alpha and Delta variants with potency drastically increased over tens of folds. This PERx-enabled covalent nanobody strategy and uncovered insights on potency increase can be valuable to developing effective therapeutics for various viral infections.

## INTRODUCTION

In December 2019 a novel coronavirus (SARS-CoV-2) caused an outbreak of coronavirus disease-19 (Covid-19) pandemic,^1^ which has been ongoing for over 2 years and resulted in an unprecedented burden to public health globally. In addition to vaccines,^2^ incessant efforts have been spent on developing drugs to inhibit the virus as prophylactics and treatment. Since cell entry of SARS-CoV-2 depends on the binding of the viral Spike protein to the human cellular angiotensin-converting enzyme 2 (ACE2) receptor,^3, 4^ various types of reagents have been developed to block the Spike-ACE2 interaction to neutralize SARS-CoV-2. ^5 6 7 8 9 10 11 12^ Despite the exciting progress, complete inhibition of SARS-CoV-2 infection remains challenging. Biologics such as protein-based drugs exert their function through non-covalent interactions, which are reversible to allow drug dissociation such that the unblocked virus can re-access and infect cells. In addition, rapid evolving of the SARS-CoV-2 RNA genome has led to variants that escape neutralization by human immune system or various noncovalent reagents.^13 14 15^ Covalent small molecule drugs have been shown to possess enhanced potency, prolonged duration of action, and ability to mitigate drug resistance.^16 17^ We thus envisioned that protein drugs able to bind the virus irreversibly in covalent mode would be highly desirable to attain potent and complete inhibition of viral infection, as well as to minimize viral escape through mutation. However, whether and how covalent protein drugs can increase potency in neutralizing viral infection await exploration.

Natural proteins generally lack the ability to bind with target covalently.^18^ To break this natural barrier, we recently reported a Proximity-Enabled Reactive therapeutics (PERx) strategy to generate covalent protein drugs.^19 20^ A latent bioreactive unnatural amino acid (Uaa) is incorporated into the protein drug through genetic code expansion,^21^ which reacts with a natural residue on the target only upon drug-target binding, realizing specific cross-linking of the drug to the target covalently. We have demonstrated that a covalent PD-1 drug efficiently inhibits tumor growth in mice, with therapeutic efficacy superior to that of an FDA-approved antibody.^19^ This initial success was achieved on suppression of tumor growth, which is a relatively long time process taking days to weeks. It remains to be established whether PERx can be generally applicable to other proteins for enhanced efficacy and to acute processes requiring fast and prompt reaction.

Here we developed covalent nanobodies to irreversibly inhibit the infection of both SARS-CoV-2 and its variants via the PERx principle. To cope with acute viral infection, a novel latent bioreactive Uaa, fluorine substituted fluorosulfate-L-tyrosine (FFY), was designed and genetically encoded to accelerate the PERx reaction rate by 2.4 fold over the original FSY, enabling fast cross-linking within 10 min. Mechanistic insight was uncovered on how to translate the increase in antagonizing Spike-ACE2 interaction into more potent virus neutralization. Consequently, FFY-based nanobodies exhibited drastic potency increase in neutralizing SARS-CoV-2 (41-fold) as well as its Alpha (23-fold) and Delta (39-fold) variants. Using PERx strategy, covalent soluble human ACE2 was also generated to bind to the Spike protein irreversibly. This PERx-based covalent nanobody strategy may provide a new route to developing effective therapeutics for viral infections.

## RESULTS

### Covalent nanobodies to neutralize SARS-CoV-2 via PERx

Nanobodies (single-domain antibodies) are generally heat stable, easier to produce in bacteria, have small size (∼15 kDa) to increase binding density on virus for efficient blockage,^8^ and can be humanized to minimize potential immunogenicity. Multiple groups have selected nanobodies binding to the Spike protein with high affinities, showing promising inhibition against SARS-CoV-2 infection of Vero cells or ACE2-expressing HEK-293 cells.^22, 23 7^ Our strategy was to genetically incorporate a latent bioreactive unnatural amino acid (Uaa) into the nanobody that specifically binds to the receptor binding domain (RBD) of the SARS-CoV-2 viral Spike protein (**Figure 1a**). Upon binding of the nanobody with the Spike RBD, the latent bioreactive Uaa would be brought into close proximity to a target residue of the Spike RBD, which enables the Uaa to react with the target residue specifically, irreversibly cross-linking the nanobody with the Spike RBD. The bound nanobody would prevent the Spike RBD interact with human ACE2 receptor, blocking viral infection. The resultant covalent nanobody would thus function as proximity-enabled reactive therapeutics (PERx). Compared with the conventional nanobodies, which bind in non-covalent mode and are in dynamic association and dissociation with Spike RBD, covalent nanobodies would permanently bind to Spike RBD and neutralize the virus with enhanced potency (**Figure 1b**).

**Figure 1.**
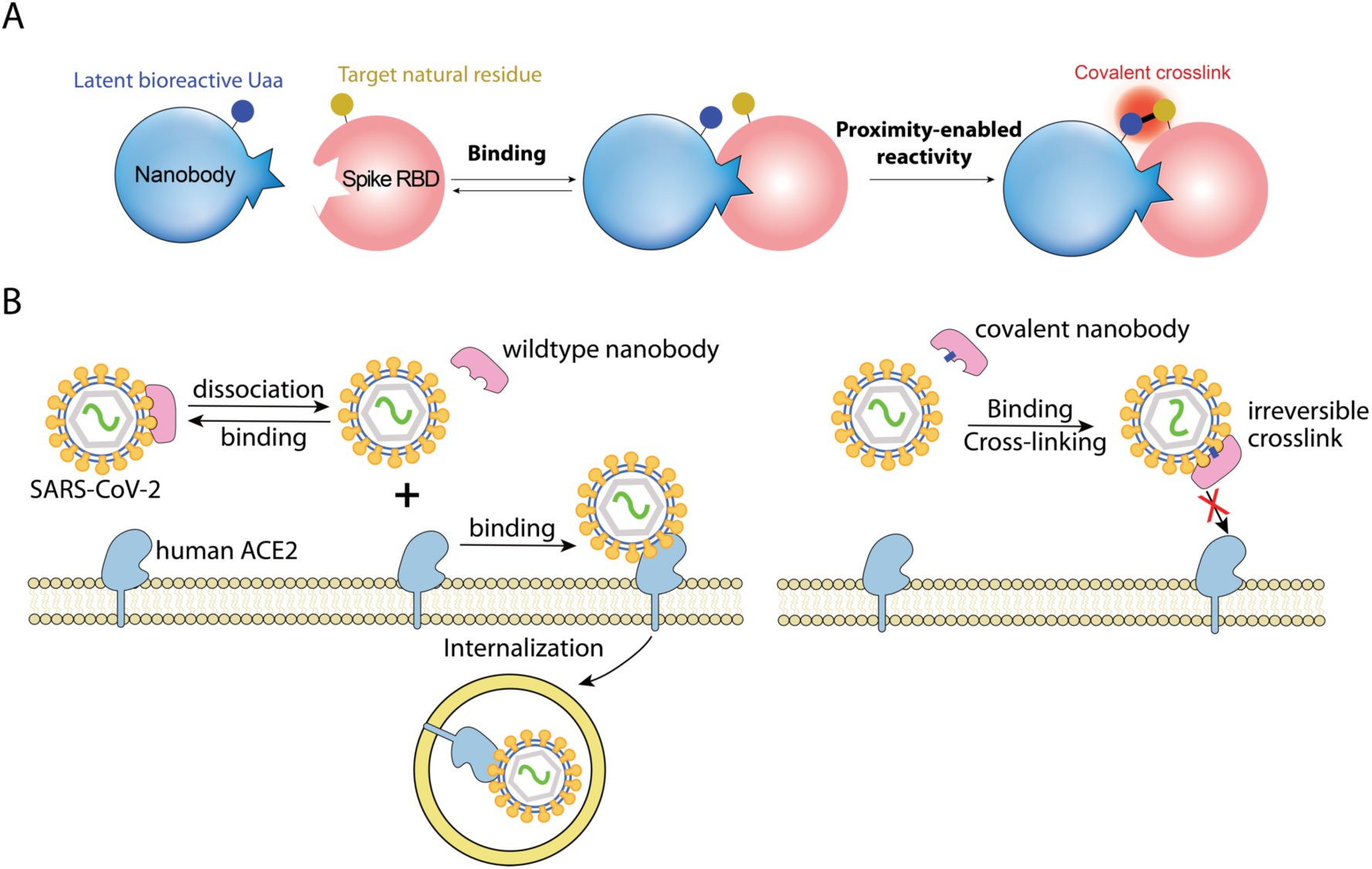
Covalent nanobody irreversibly inhibits SARS-Cov-2 infection via PERx. (**A**) The PERx principle of developing covalent nanobody to irreversibly bind with the Spike RBD. Only upon nanobody binding with the Spike RBD, the latent bioreactive Uaa reacts with the target natural residue, covalently cross-linking the nanobody with the Spike RBD. (**B**) While conventional nanobody can dissociate from the Spike RBD, covalent nanobody binds with the Spike RBD irreversibly, permanently preventing viral binding with ACE2 receptor and thus blocking infection more effectively.

### Genetically encoding FSY to generate covalent nanobodies that bind to Spike RBD irreversibly *in vitro*

We initially chose to use the latent bioreactive Uaa FSY for its stability and ability to react with Tyr, His, or Lys via proximity-enabled sulfur fluoride exchange (SuFEx) reaction under biocompatible and cellular conditions (**Figure 2A**).^24^ On the basis of the crystal structure of human SARS-CoV-2 Spike RBD in complex with nanobody H11-D4,^25^ nanobody MR17-K99Y,^23^ or nanobody SR4,^23^ we decided to incorporate FSY at site R27, S30, E100, W112, D115, or Y116 of nanobody H11-D4 (**Figure 2B**), site Y99 or D101 of nanobody MR17-K99Y (**Figure 2C**), and site Y37, H54 or S57 of nanobody SR4 (**Figure 2D**), respectively. Western blot analysis of lysates of cells expressing these mutant nanobody genes and the tRNA^Pyl^/FSYRS pair^24^ confirmed that FSY was successfully incorporated into the nanobodies in the presence of FSY (**Figure S1**). WT and FSY-incorporated nanobodies were purified with affinity chromatography. Mass spectrometric analysis of the intact SR4(57FSY) protein confirmed that FSY was incorporated into SR4 at site 57 with high fidelity (**Figure 2E**).

**Figure 2.**
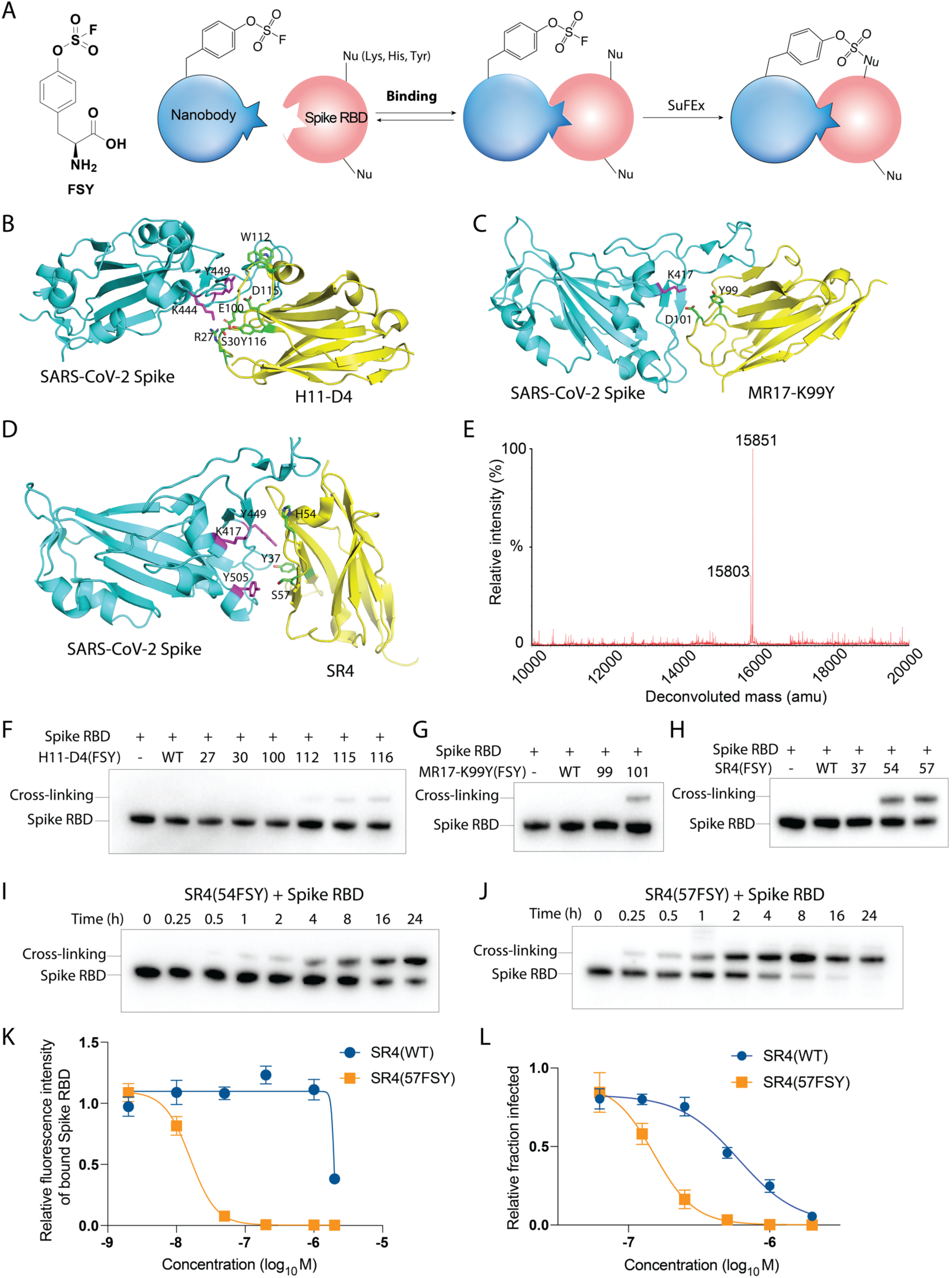
FSY-incorporated covalent nanobody cross-linked the Spike RBD and more potently inhibited Spike RBD binding with cell surface ACE2, as well as pseudoviral infection. (**A**) Structure of FSY and its proximity-enabled SuFEx reaction with Lys, His, or Tyr for PERx. (**B-D**) The crystal structure of nanobody in complex with SARS-CoV-2 Spike RBD. (B) Nanobody H11-D4 (PDB: 6YZ5); (C) Nanobody MR17-K99Y (PDB: 7CAN); (D) Nanobody SR4 (PDB: 7C8V). Sites selected for FSY incorporation in the nanobody and intended target residues of the Spike RBD are shown in green and magenta stick, respectively. (**E**) ESI-MS spectrum of the intact SR4(57FSY) confirming FSY incorporation. Expected 15802.5 Da; Observed 15803 Da. The peak observed at 15851 Da corresponded to SR4(57FSY) with Cysteine oxidation. (**F-H**) Cross-linking of the Spike RBD with wildtype nanobody H11-D4 (F), MR17-K99Y (G), SR4 (H) and their FSY mutants at 37 °C for 12 h. Mouse Fc tag appended at the C-terminus of the Spike RBD was detected in the Western blot. (**I-J**) Western blot analysis of 5 μM SR4(54FSY) or SR4(57FSY) cross-linking with 0.5 μM Spike RBD at indicated time points. (**K**) Inhibition curve of the Spike RBD-mFc binding to 293T-ACE2 cells by SR4(WT) or SR4(57FSY). SR4(WT) or SR4(57FSY) (in final conc. 2, 1, 0.2, 0.05, 0.01, and 0.002 μM) were used to inhibit 10 nM Spike RBD-mFc binding to 293T-ACE2 cells. Fluorescence intensity representing the Spike RBD-mFc bound to 293T-ACE2 cells was measured using FITC-labeled mFc antibody by flow cytometry. Error bars represent s.d., n = 3 biological replicates. (**L**) Inhibition of SARS-CoV-2 pseudovirus infection of 293T-ACE2 cells by SR4(WT) or SR4(57FSY). The percentage of GFP positive cells, the indicator for infection, was measured with flow cytometry. The normalized infection in y-axis was calculated as (the percentage of GFP positive cells infected by nanobody-treated pseudovirus) / (the percentage of GFP positive cells infected by pseudovirus only). Error bars represent s.d., n = 3 biological replicates.

To test if FSY-incorporated nanobodies could bind to the Spike RBD covalently *in vitro*, we incubated 2.5 μM WT nanobodies or their FSY mutants with 0.5 μM Spike RBD at 37 °C for 12 h, and then performed Western blot analysis under denatured conditions (**Figure 2F-H**). As expected, all three WT nanobodies did not form a covalent complex with the Spike RBD. A stable covalent complex was detected for H11-D4(115FSY) and H11-D4(116FSY) with low cross-linking efficiencies (**Figure 2F**), for MR17-K99Y(101FSY) with 10.5 % cross-linking efficiency (**Figure 2G**), for SR4(54FSY) with 28.3 % cross-linking efficiency, and SR4(57FSY) with 41.3 % cross-linking efficiency (**Figure 2H**). We then further studied time-dependent cross-linking of the Spike RBD with SR4(54FSY) or SR4(57FSY) nanobodies, which showed SR4(57FSY) was more efficient (**Figure 2I-J**). SR4(57FSY) was thus used in subsequent experiments.

### SR4(57FSY) inhibits binding of Spike RBD to cell surface ACE2 receptor

We next tested the efficacy of SR4 nanobodies to inhibit the binding of the Spike RBD-mFc (a mouse Fc tag appended at the C-terminus of Spike RBD) to 293T-ACE2 cells, a HEK293T cell line stably expressing human ACE2 protein on cell surface. Different concentrations of SR4(WT) or SR4(57FSY) were individually incubated with the Spike RBD-mFc at 37 °C for 12 h to allow cross-linking, followed by incubation with 293T-ACE2 cells for 1 h. After incubation, cells were stained with FITC labeled anti-mFc and analyzed with flow cytometry (**Figure 2K**). As expected, SR4(WT) bound to the Spike RBD reversibly such that the Spike RBD could still bind to ACE2 on the cell surface, resulting in an IC_50_ of 1980 nM. In contrast, the covalent SR4(57FSY) showed highly efficient blocking of the Spike RBD binding to 293T-ACE2 cells with an IC_50_ of 15.8 nM (**Figure 2K**). The IC_50_ of SR4(57FSY) was 125-fold lower than that of SR4(WT), demonstrating the drastic improvement of the covalent nanobody in inhibiting the binding of Spike RBD to cell surface ACE2 receptor.

### SR4(57FSY) neutralizes pseudovirus infection of 293T-ACE2 cells

We next tested the neutralization activity of SR4(WT) or SR4(57FSY) nanobody against SARS-CoV-2 spike pseudotyped lentivirus using a widely adopted protocol.^26 27^ SARS-CoV-2 reporter virus particles display antigenically correct Spike protein on a heterologous virus core and carry a modified genome that expresses a convenient GFP reporter gene, which is integrated and expressed upon successful viral entry into cells harboring the ACE2 receptor. Briefly, the pseudoviruses were incubated with various concentrations of SR4(WT) or SR4(57FSY) at 37 °C for 1 h, and subsequently used to infect 293T-ACE2 cells for 48 h. Cells were then analyzed by flow cytometry for GFP signal to determine the percentage of infected cells. The IC_50_ for SR4(WT) was measured to be 602.8 nM (**Figure 2L**), close to the literature reported 390 nM,^23^ whereas the IC_50_ for SR4(57FSY) was measured to be 151.6 nM. This 4-fold decrease of IC_50_ demonstrated that the covalent SR4(57FSY) was more potent in inhibiting pseudovirus infection of ACE2-expressing human cells than SR4(WT), but the enhancement was moderate.

We further tested the neutralization ability of SR4(WT) or SR4(57FSY) nanobody against authentic SARS-Cov-2 infection of ACE2 expressing human cells. Surprisingly, no significant enhancement of SR4(57FSY) over SR4(WT) was measured (Data not shown). The lack of enhancement in inhibiting authentic virus infection seemed to be discrepant with the fact that SR4(57FSY) drastically enhanced inhibition of the Spike RBD binding with cell surface ACE2 receptor (125 fold). We reasoned that the discrepancy could be accounted for by how SR4 accesses and binds to the Spike RBD on the live SARS-CoV-2 virus, which may be different from the purified Spike RBD in isolation. In fact, the Spike RBD in SARS-Cov-2 virus exchanges between the active up state and the inactive down state.^28,29^ To achieve potent inhibition of viral infection, binding and locking the Spike RBD in the inactive down state could be more effective.^7^ However, the crystal structure of SR4 in complex with RBD indicates that SR4 binds with the up state of RBD instead.^23^ We therefore sought to covalently engineering a nanobody that could bind the Spike RBD in the inactive down state.

### Generate covalent nanobodies from mNb6 that binds the down-state of RBD

Nanobody mNb6 was isolated through screening a library against the Spike ectodomain stabilized in the prefusion conformation, and thus bound the Spike RBD in the down state.^7^ To identify which sites of mNb6 would allow covalent crosslink of the Spike RBD, we incorporated FSY individually at 30 different sites located at the three complementarity-determining regions (CDR) of mNb6 (**Figure 3A**). Although the crystal structure of mNb6 in complex with RBD is available,^7^ we performed this comprehensive site screening without inspecting the structure to show that for proteins with well-defined regions, such as the nanobody, one could readily determine the appropriate sites for FSY cross-linking without sophisticated high-throughput screening or detailed structures. We incubated 2 μM mNb6(WT) and its FSY mutant proteins with 0.5 μM Spike RBD in PBS (pH 7.4) at 37 °C for 12 h, followed by Western blot analysis (**Figure 3B-D**). When FSY was incorporated at site 27 in CDR1, site 55 in CDR2, or sites 102-108 in CDR3, a covalent complex of mNb6 with the Spike RBD was detected, indicating that multiple sites in mNb6 allowed covalent cross-linking with the Spike RBD, which may not be obvious by only inspecting the crystal structures. We then performed kinetic study on the four more efficient sites on CDR3 (**Figure 3E**), and found that mNb6(108FSY) showed the fastest cross-linking rate. We thus chose site 108 to incorporate Uaa for subsequent experiments.

**Figure 3.**
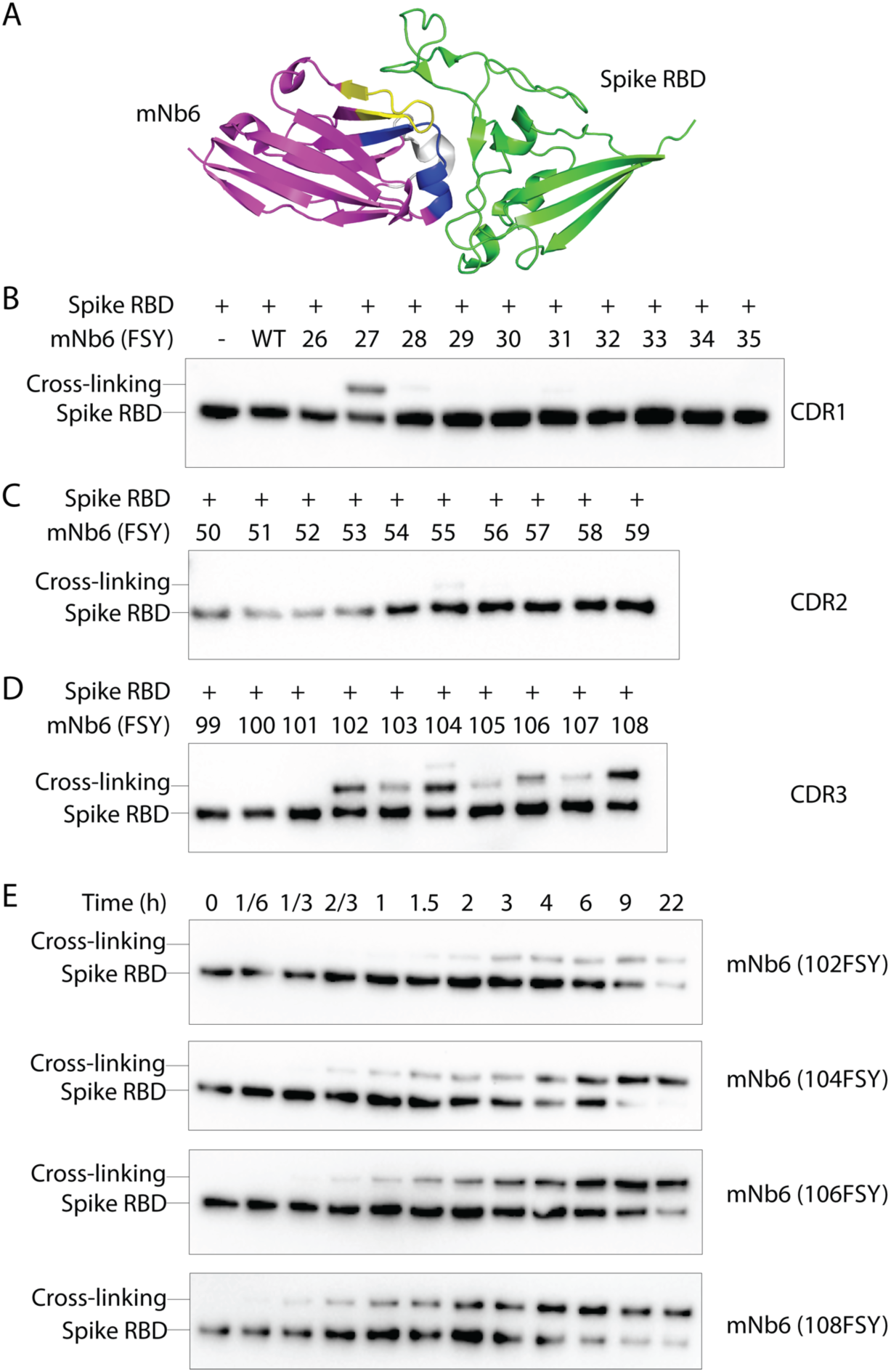
Covalent nanobodies based on mNb6 that binds the down-state of Spike RBD. (**A**) Crystal structure of Spike RBD binding with mNb6 (PDB: 7KKL). FSY was individually incorporated at 30 different positions located at the CDR1 (blue), CDR2 (yellow), and CDR3 (gray) region of the nanobody without referring to the specific structure of mNb6. (**B-D**) Western blot analysis of cross-linking between the Spike RBD and WT mNb6 or FSY mutants located at 3 CDRs of the nanobody, respectively. (**E**) Western blot analysis of mNb6(FSY) mutants (5 μM) cross-linking with the Spike RBD (0.5 μM) at indicated time points.

### Fluorine substituted fluorosulfate-L-tyrosine (FFY) as a novel latent bioreactive Uaa to accelerate PERx reaction rate

The potency of covalent protein drug would rely on the rate and extent of latent bioreactive Uaa to form a covalent bond with the target residue of the target protein. As protein interactions are dynamic, a fast reaction rate would be essential to ensure covalent bond formation before protein dissociation. Given certain contact time, the reaction extent increases with faster reaction rate, which can be critical to achieve complete inhibition of viral infection. Introducing electron-withdrawing groups on the aromatic ring has been reported to increase the SuFEx rates.^30^ We thus envisioned that adding electron-withdrawing substituents to FSY would accelerate its proximity-enabled reaction rate when used in PERx.

As a result, we designed and evaluated a fluorine substituted fluorosulfate-L-tyrosine (FFY, **Figure 4A**). FFY was synthesized using [4-(acetylamino)phenyl]imidodisulfuryl difluoride (AISF),^31^ followed with the deprotection of the Boc protecting group using hydrogen chloride. As FFY and FSY have similar structures, we reasoned that FSYRS, a pyrrolysyl-tRNA synthetase (PylRS) mutant we previously evolved to incorporate FSY,^24^ should be able to incorporate FFY into proteins as well. To test this idea, the enhanced green fluorescent protein (EGFP) gene containing a TAG codon at permissive site 182 was co-expressed with genes for tRNA^Pyl^/FSYRS in *E. coli*. In the absence of FFY, no obvious fluorescence was detected; in the presence of FFY, fluorescence intensity was measured to increase with FFY concentration (**Figure 4B**), suggesting FFY incorporation into EGFP. We also co-expressed mNb6(108TAG) with tRNA^Pyl^/FSYRS in *E. coli*, and observed that full-length mNb6 was produced in the presence of 2 mM FFY or 1 mM FSY (**Figure 4C**). mNb6(WT), mNb6 (108FSY), and mNb6(108FFY) proteins were purified with Ni^2+^ affinity chromatography. Mass spectrometric analysis of the intact protein confirmed that FFY was incorporated into mNb6 at site 108 in high fidelity. A major peak observed at 13721 Da corresponds to intact mNb6(108FFY) (**Figure 4D**, expected 13720.7 Da). A minor peak observed at 13702 Da corresponding to mNb6(108FFY) lacking F, suggesting a slight F elimination during mass spectrometric measurement.^24^ We also verified mNb6(WT) and mNb6(108FSY) via mass spectrometric analysis of the intact proteins (**Figure 4E** and **Figure S2**).

**Figure 4.**
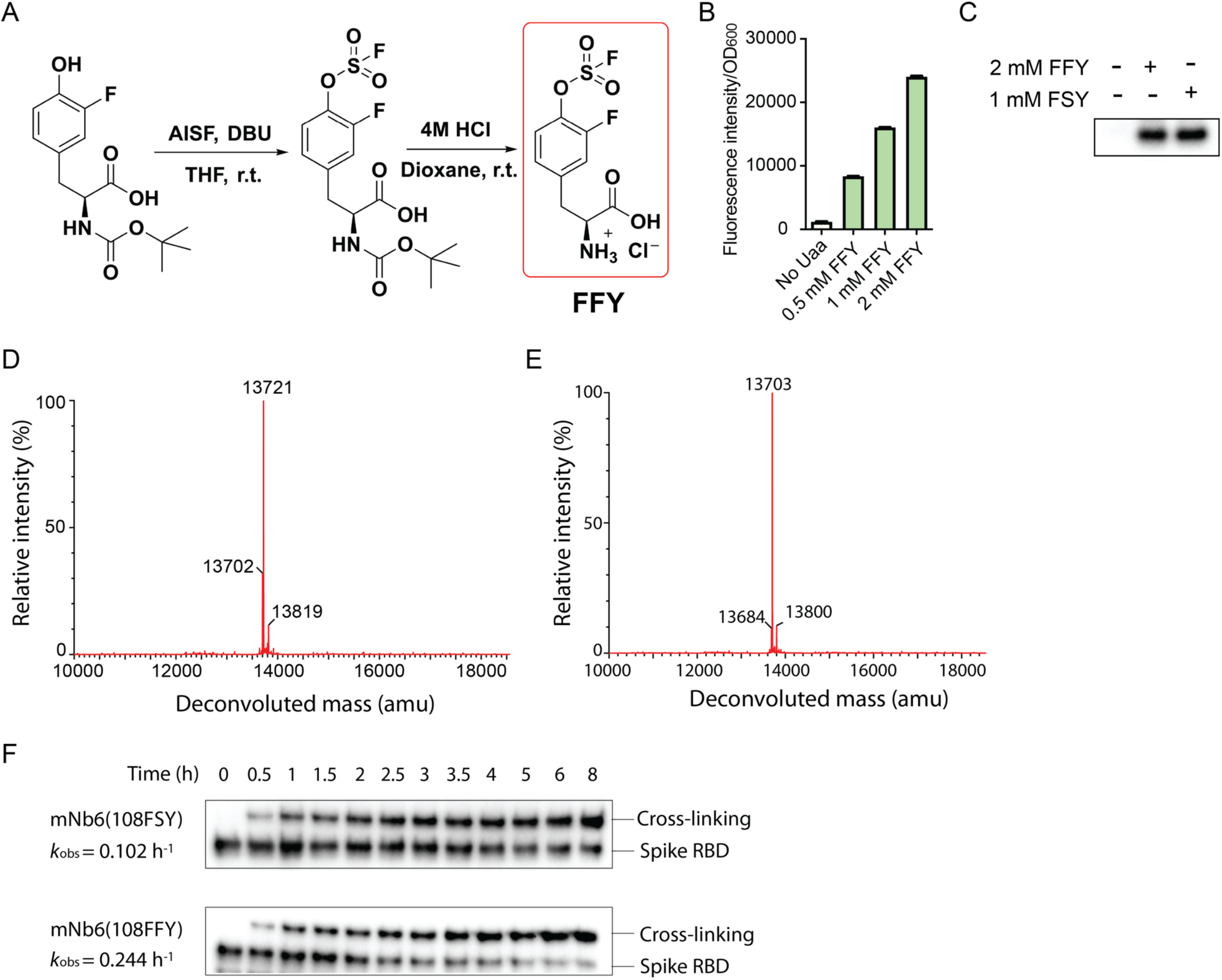
Design and genetically encode FFY to accelerate PERx reaction rate. (**A**) Chemical synthesis of FFY. (**B**) Genetic incorporation of FFY into EGFP at site 182 in *E. coli* using the tRNA^Pyl^/FSYRS pair. Fluorescence intensity of cells was measured and normalized by OD_600_. (**C**) Genetic incorporation of FFY or FSY into mNb6 at site 108 in *E. coli* using the tRNA^Pyl^/FSYRS pair. Cell lysates were analyzed with Western blot using an anti-Hisx6 antibody. (**D**) Mass spectrometric analysis of the intact mNb6(108FFY). Expected 13720.7 Da, observed 13721 Da. (**E**) Mass spectrometric analysis of the intact mNb6(108FSY). Expected 13702.7 Da, observed 13703 Da. The minor peak at 13684 Da corresponding to mNb6(108FSY) lacking F, suggesting slight F elimination during MS measurement. (**F**) Western blot analysis of cross-linking kinetics indicates that FFY increased the reaction rate to 2.4 fold over FSY. The Spike RBD (0.5 μM) was incubated with 5 μM mNb6(108FFY) or mNb6(108FSY) in PBS (pH = 7.4) at 37 °C for indicated time.

We next tested if FFY could improve reaction kinetics over FSY. The Spike RBD (0.5 μM) was incubated with 5 μM mNb6(108FSY) or mNb6(108FFY) in PBS (pH 7.4) at 37 °C for different duration, followed by Western blot analysis of the cross-linking. As shown in **Figure 4F**, mNb6(108FFY) showed evidently faster rate than mNb6(108FSY) in cross-linking the Spike RBD. Covalent cross-linking could be robustly detected within 10 min for mNb6(108FFY) (**Figure S3**), while mNb6(108FSY) took over 20 min. The apparent first-order constant *k*_obs_ was 0.244 ± 0.031 h^-1^ (n = 3) for mNb6(108FFY) and 0.102 ± 0.007 h^-1^ (n = 3) for mNb6(108FSY) (**Figure 4F**, quantification in **Figure S4**). Thus, FFY increased the PERx reaction rate over FSY in the mNb6 system to 240%, so we used mNb6(108FFY) for subsequent viral inhibition tests.

### mNb6(108FFY) potently neutralizes both pseudovirus and authentic SARS-Cov-2 infection

With FFY’s fast kinetics and mNb6 binding of the down state of Spike RBD, we tested the efficacy of mNb6(108FFY) in neutralizing SARS-CoV-2 spike pseudotyped lentivirus infection. Pseudovirus was incubated with various concentrations of mNb6(108FFY) or mNb6(WT) for 1 h at 37 °C, and then the nanobody-pseudovirus mixture was used to infect 293T-ACE2 cells for 48 h at 37 °C. The percentage of cell infection was determined by measuring GFP signal via flow cytometry. The IC_50_ of mNb6(WT) was measured to be 35.7 nM. In contrast, mNb6(108FFY) showed an IC_50_ of 1.0 nM, exhibiting a marked 36-fold improvement in potency (**Figure 5A**). To find out if FFY incorporation impacted the binding affinity of mNb6, we measured the binding affinity of mNb6(WT) or mNb6(108FFY) toward the Spike RBD using biolayer interferometry (BLI). The *K*_D_ was measured to be 0.99 nM for both mNb6(WT) and mNb6(108FFY) (**Figure 5B-C**), indicating that FFY incorporation did not significantly affect mNb6 binding affinity. Therefore, the drastically elevated potency of mNb6(108FFY) over mNb6(WT) in neutralizing pseudovirus infection should be attributed not to affinity difference but to the effect of covalent binding.

**Figure 5.**
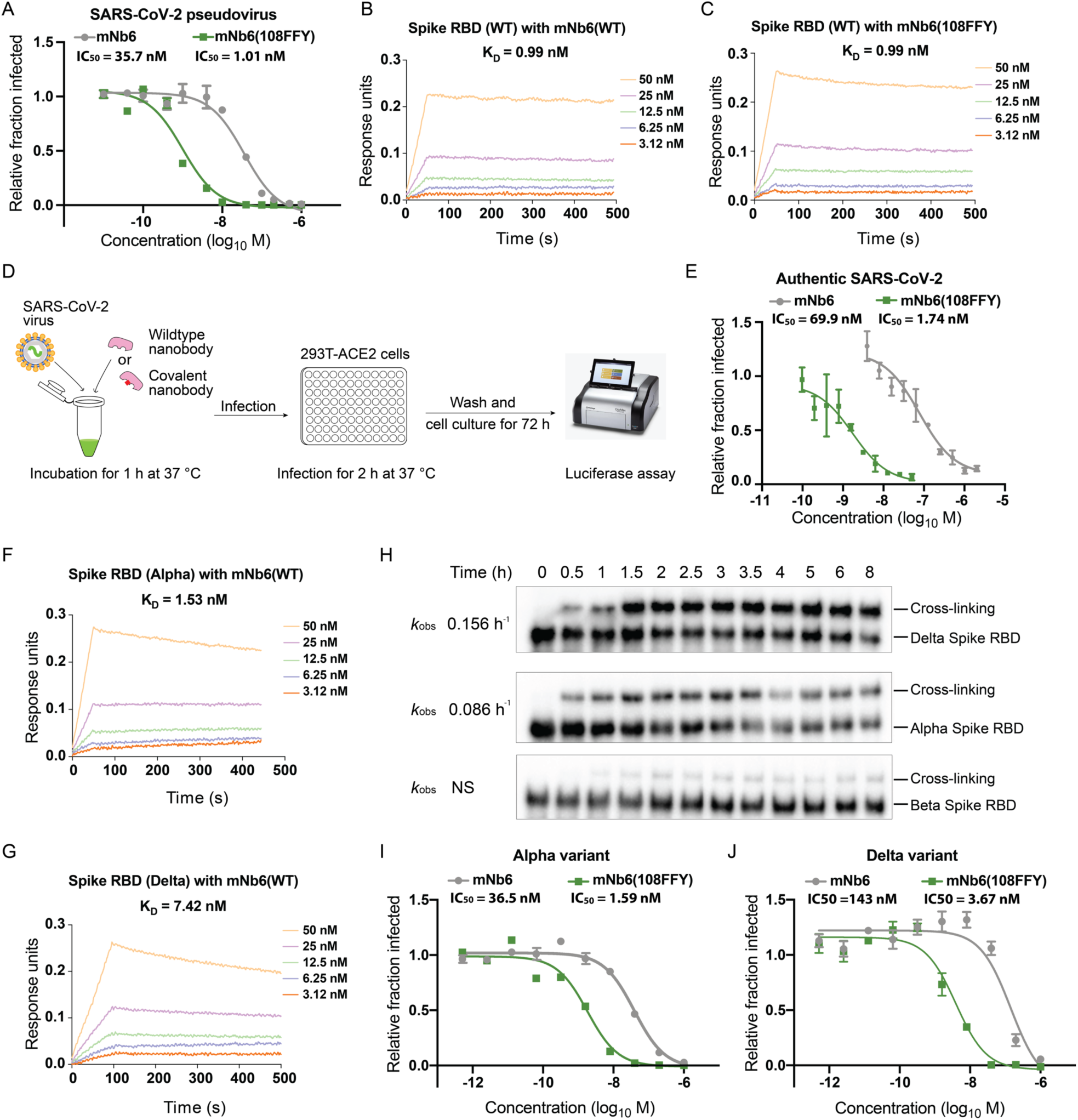
mNb6(108FFY) neutralizes SARS-Cov-2 and certain variants with markedly increased potency over mNb6(WT). (**A**) mNb6(108FFY) showed 36-fold increase in potency than mNb6(WT) in inhibiting SARS-CoV-2 pseudovirus infection. (**B-C**) BLI of mNb6(WT) or mNb6(108FFY) binding to the Spike RBD, showing no difference in *K*_D_. (**D**) Scheme showing procedures for neutralizing authentic SARS-CoV-2 with WT or covalent nanobody. (**E**) mNb6(108FFY) showed 41-fold increase in potency than mNb6(WT) in inhibiting authentic SARS-CoV-2 infection. **(F-G**) BLI of mNb6(WT) binding to the RBD of Alpha (F) or Delta (G) variant of SARS-Cov-2. (**H**) Cross-linking of mNb6(108FFY) with the Spike RBD of SARS-CoV-2 variants at indicated incubation time *in vitro*. (**I**) mNb6(108FFY) showed 23-fold increase in potency than mNb6(WT) in inhibiting the Alpha variant pseudovirus infection. (**J**) mNb6(108FFY) showed 39-fold increase in potency than mNb6(WT) in inhibiting the Delta variant pseudovirus infection. For all pseudovirus and authentic SARS-CoV-2 inhibition experiments in (A), (E), (I), and (J), n = 3 biological repeats; error bars represent s.d..

Encouraged by the highly potent effect on SARS-CoV-2 pseudovirus, we further tested the neutralizing efficacy of mNb6(108FFY) on authentic SARS-CoV-2 virus (**Figure 5D**). Nanoluciferase-encoding SARS-CoV-2 was incubated with the nanobodies at 37 °C for 1 h, and the virus-nanobody mixtures were then added to 293T-ACE2 cells for 2 h to allow infection. At 72 h post-infection, luciferase signal in cells was measured to determine the level of infection. As shown in **Figure 5E**, the IC_50_ of mNb6(WT) in inhibiting authentic SARS-CoV-2 was 69.9 nM whereas the IC_50_ measured for mNb6(108FFY) was 1.7 nM, indicating a drastic 41-fold improvement in potency.

### mNb6(108FFY) covalently binds and potently inhibits certain SARS-CoV-2 variants

Various mutated SARS-CoV-2 strains have emerged during the pandemic. The B.1.1.7 variant (Alpha), which possesses the N501Y mutation, shows a stronger interaction with ACE2 and a faster spreading rate;^32 33^ the B.1.351 variant (Beta), which possesses K417N, E484K, and N501Y mutations on the RBD, has decreased affinity towards neutralization antibodies and can lower the effectiveness of current vaccine.^34 13 35 14^ More recently, the B.1.617.2 variant (Delta), which has the L452R and T478K mutations on the RBD, has become the dominant strains in many countries. Delta variant has increased transmissibility and is less sensitive to monoclonal antibodies and neutralizing antibodies from recovered individuals, as well as vaccine-elicited antibodies.^15 36 37^

We first tested whether these variants would escape from mNb6(WT) binding. BLI measurement showed that mNb6 had a decreased binding affinity towards the RBD of the Alpha and Delta variants. The *K*_D_ of mNb6(WT) with the Alpha and Delta RBD was 1.53 and 7.42 nM, respectively (**Figure 5F-G**), higher than 0.99 nM, the *K*_D_ with the WT RBD. In contrast, mNb6(WT) failed to show significant binding with the Beta RBD at the highest concentration tested (50 nM) (**Figure S5**). We next determined whether mNb6(108FFY) could crosslink the RBD of various variants. Western blot analysis indicated that mNb6(108FFY) efficiently cross-linked with the Delta and Alpha RBD (**Figure 5H**, quantification in **Figure S6**), although the cross-linking rate (*k*_obs_ = 0.156 ± 0.036 h^-1^ for Delta and 0.086 ± 0.008 h^-1^ for Alpha) was slightly slower than that with the WT RBD (*k*_obs_ = 0.244 ± 0.031 h^-1^), consistent with the decreased affinity towards these variants. For the Beta RBD, although the affinity was dramatically reduced, mNb6(108FFY) was still able to crosslink it albeit in low efficiency (∼ 5%) (**Figure 5H**). These results demonstrate that the covalent mNb6(108FFY) was capable of cross-linking the Spike RBD of the SARS-Cov-2 variants, as long as the variant did not completely abolish binding with mNb6.

We further assessed the efficacy of mNb6(108FFY) in neutralizing pseudovirus of SARS-CoV-2 variants. The IC_50_ of mNb6(WT) in inhibiting the Alpha variant pseudovirus was 36.5 nM (**Figure 5I**) whereas mNb6(108FFY) had an IC_50_ of 1.6 nM, showing a 23-fold increase in potency. For the quickly spreading Delta variant, mNb6(WT) inhibited the pseudovirus with an IC_50_ of 143 nM (**Figure 5J**) whereas mNb6(108FFY) was able to neutralize the Delta variant pseudovirus with an IC_50_ of 3.7 nM, achieving a 39-fold improvement in potency. In comparison with the WT SARS-CoV-2 pseudovirus, mNb6(WT) inhibited the Delta variant pseudovirus with a much higher IC_50_ (143 nM vs 35.7 nM), which is consistent with the decreased affinity of mNb6(WT) towards the Delta RBD than the WT one (7.42 nM vs 0.97 nM). However, despite the marked decrease in binding affinity of mNb6(WT) towards Delta RBD, mNb6(108FFY) still inhibited the Delta variant pseudovirus effectively. In short, the covalent mNb6(108FFY) nanobody also potently inhibited the Alpha and Delta variant pseudoviruses, with respective 23-and 39-fold increase in potency than the mNb6(WT) nanobody.

### Engineering human ACE2 receptor into an irreversible covalent binder for SARS-CoV-2

In addition to artificial nanobodies specific for the Spike RBD, we also sought to engineer the native receptor, human ACE2, into a covalent binder to block the interaction of SARS-CoV-2 Spike with human ACE2. Using a soluble ACE2 receptor binding to the viral Spike protein, thereby neutralizing SARS-CoV-2, is an attractive strategy. The Spike protein of SARS-CoV-2 binds to the ACE2 receptor with a *K*_D_ of 4.7 nM,^38^ comparable to affinities of mAbs. In addition, ACE2 administration could additionally treat pneumonia caused by SARS-CoV-2. Coronavirus binding leads to ACE2 protein shedding and downregulation, which induces pulmonary edema and acute respiratory distress syndrome (ARDS). Administration of recombinant human ACE2 improves acute lung injury and reduces ARDS in preclinical studies.^39,40^ Moreover, recombinant human ACE2 is safe and well tolerated by patients in phase II trial,^41,42^ and small levels of soluble ACE2 are secreted and circulate in human body.^43^ More importantly, the soluble ACE2 therapy is expected to have broad coverage as SARS-CoV-2 cannot escape ACE2 neutralization due to its dependence on the same protein for cell entry. Any mutation of SARS-CoV-2 reducing its affinity for the ACE2-based therapeutics will render the virus less pathogenic.

To this end, we explored if the soluble human ACE2 could be engineered into a covalent binder for SARS-CoV-2 Spike protein. On the basis of the crystal structure of the SARS-CoV-2 Spike RBD in complex with human ACE2 (**Figure 6A**),^38^ we decided to incorporate FSY at site D30, H34, E37, D38, Q42 or Y83 to target the proximal nucleophilic amino acid residues on the Spike RBD. These FSY-incorporated ACE2 mutant proteins were expressed and purified from HEK293 cells and individually incubated with the Spike RBD followed by Western blot analysis (**Figure 6B**). Cross-linking of ACE2(FSY) with the Spike RBD was detected when FSY was incorporated at sites 34, 37, or 42. The ACE2 (34FSY) mutant exhibited the highest cross-linking efficiency (28.3%). The cross-linking kinetic experiments of ACE2(FSY) with the Spike RBD showed that cross-linking was detectable at 1 h and increased with incubation time (**Figure 6C**). Furthermore, we analyzed the cross-linking product of ACE2(34FSY) with the Spike RBD using high-resolution mass spectrometry, which clearly indicated that FSY of ACE2(34FSY) specifically reacted with K417 of the Spike RBD (**Figure S7**), as designed based on the crystal structure. Together, these results showed that ACE2(34FSY) covalently bound to the Spike RBD.

**Figure 6.**
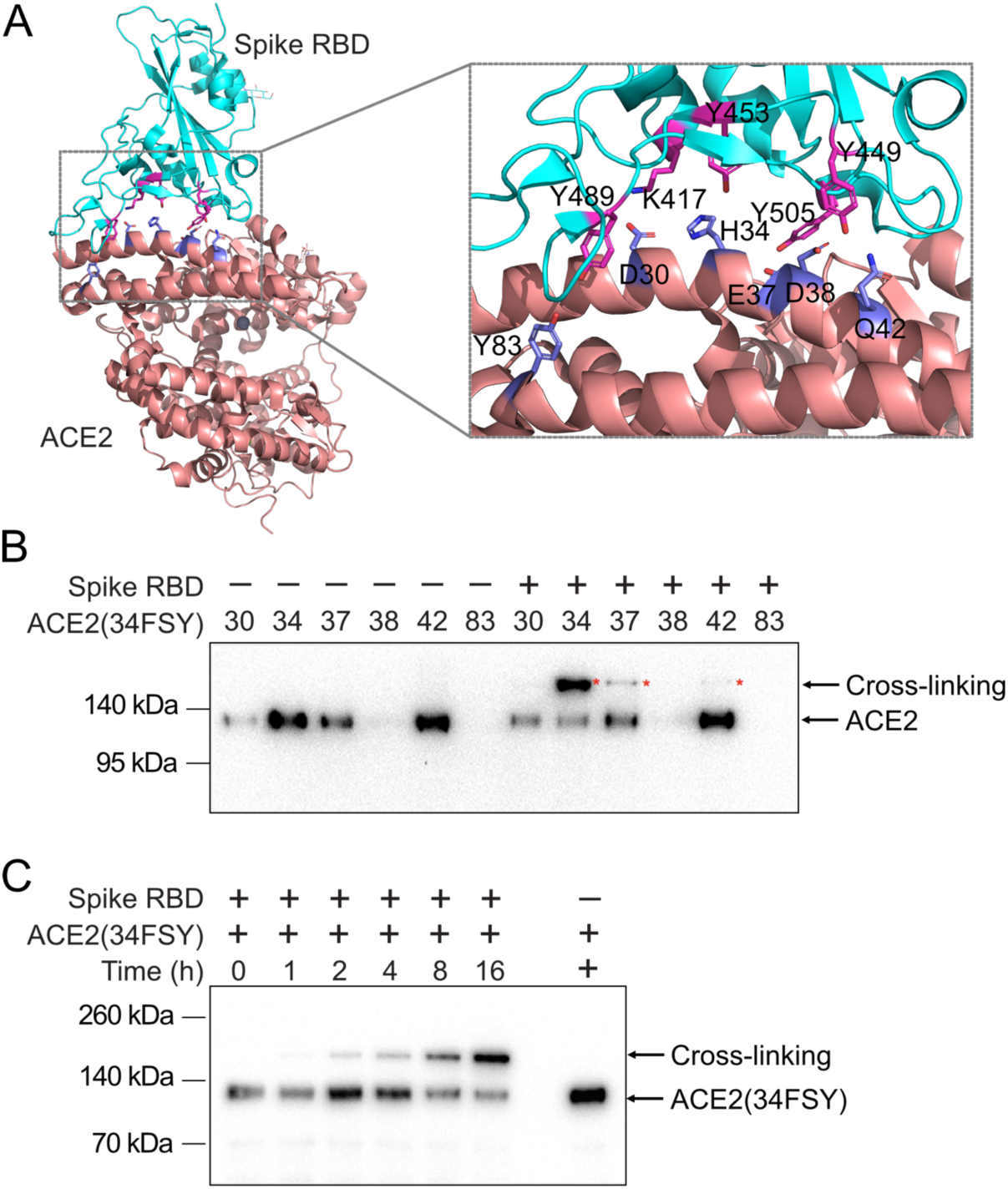
Engineering soluble human ACE2 to covalently bind to the SARS-CoV-2 Spike protein. (**A**) The crystal structure of SARS-CoV-2 Spike RBD bound with human ACE2 (PDB: 6M0J). Sites on ACE2 selected to incorporate FSY are shown in blue stick, and target residues on the Spike RBD are shown in magenta stick. (**B**) Western blot analysis of cross-linking of the Spike RBD with soluble human ACE2 and its FSY mutants at 37 °C. (**C**) Western blot analysis of ACE2(34FSY) (0.16 μM) cross-linking with the Spike RBD (0.57 μM) at indicated time duration. For both (B) and (C), an antibody against Hisx6 tag at the C-terminus of ACE2(FSY) was used for detection.

## DISCUSSION

In summary, through genetic incorporation of the latent bioreactive Uaas FSY and FFY into nanobodies, we generated multiple covalent nanobodies that bind with the Spike RBD of SARS-CoV-2 irreversibly via the PERx mechanism. The newly developed FFY increased the PERx reaction rate by 2.4 fold over the original FSY, and the resultant covalent nanobody mNb6(108FFY) efficiently neutralized both SARS-CoV-2 pseudovirus and authentic SARS-CoV-2, increasing the potency by a drastic 36-fold and 41-fold over the noncovalent WT mNb6, respectively. In addition, the covalent mNb6(108FFY) nanobody also potently inhibited the Alpha and Delta variant of SARS-CoV-2 pseudovirus, with a respective 23- and 39-fold increase in potency than the WT mNb6. Moreover, using a similar strategy, the soluble human ACE2 receptor was also engineered into a covalent binder that irreversibly cross-linked with the Spike RBD via PERx.

Our results demonstrate that nanobodies can be readily engineered into covalent binders through incorporating a latent bioreactive Uaa, which reacts with a natural residue of the target protein only upon nanobody-target binding, expanding the scope of PERx that we previously developed and applied to the immune-checkpoint PD-1/PD-L1.^19 20^ Sites in nanobody appropriate for FSY or FFY incorporation and cross-linking were identified either by inspecting the structure of nanobody-target complex ^44^ or simply screening all sites in nanobody’s three CDRs. Recent breakthrough in accurate prediction of protein structure and interactions have made structural information available for a broad range of proteins,^45 46^ which will greatly facilitate structure-guided site identification. On the other hand, there are about 30 total sites in nanobody’s three CDRs, screening of which using cross-linking and Western blot analysis can be completed in less than 2 weeks. We have always identified multiple sites in diverse nanobodies with successful cross-linking owing to FSY and FFY’s exceptional ability to react with multiple residues including Tyr, His, and Lys,^24^ which are often found at protein-protein interface. To react with residues at further distances, we have also developed FSK that has a longer and more flexible side chain with similar reactivity.^44^ These site identification strategies are straightforward and obviate sophisticated instrumentation such as mass spectrometry, and should be generally applicable to engineering various proteins such as antibodies, Fab, single-chain variable fragment (ScFv), affibodies, DARPins and so on, into covalent binders.

Compared with conventional noncovalent drugs, drugs working in covalent mode offer desirable features such as increased potency, prolonged duration, and possibility to prevent drug resistance, as demonstrated with covalent small molecule drugs.^16^ Here we demonstrate that the covalent nanobodies, representing covalent protein drugs, possessed similar valuable properties as well. In particular, the covalent mNb6(108FFY) drastically increased the potency over WT mNb6 to 36-fold in inhibiting SARS-CoV-2 pseudovirus and to 41-fold in inhibiting authentic SARS-CoV-2. Aside from the WT SARS-Cov-2, mNb6(108FFY) was also able to covalently crosslink and neutralize the Alpha and Delta variants of SARS-CoV-2, with potency increase of 23-and 39-fold over the WT mNb6. These potency increases were measured by incubating the nanobody with the virus for 1 h. Because the extent of covalent cross-linking increases with time, the potency of covalent nanobodies should be greater with longer incubation time.

A potential caveat is that the cross-linking rate can be negatively impacted when the binding affinity between protein drug and target decreases drastically due to target mutation, such as mNb6(108FFY) and Beta RBD. In rare cases, mutation can even occur at the cross-linking residue, which would abolish cross-linking at that site. However, as we demonstrated here and previously,^19^ _44_ PERx is a general strategy that can be applied on diverse nanobodies and each nanobody can have FFY incorporated at multiple sites to target different natural residues on Spike RBD to achieve crosslink (**Figure 3**). Therefore, other nanobodies able to bind mutated Spike RBD could be similarly engineered, and diverse covalent nanobodies could be used in combination to ensure the cross-linking and inhibition of future potential variants. When in use, the irreversible cross-linking ability together with multi-targeting of these covalent nanobodies should achieve complete viral inhibition and mechanistically prevent drug resistance. Nonetheless, animal tests and clinical trial are warranted to confirm these potential benefits of covalent protein drugs.

Fast reaction kinetics would be critical for effective inhibition of viral infection, as it enables covalent crosslinks to promptly form between the protein drug and the target within a shorter contact time, and to reach higher extent for more effective neutralization. By introducing electron withdrawing fluorine substitution we designed and genetically encoded a novel latent bioreactive Uaa FFY, which afforded a marked 2.4-fold increase in reaction rate in the cross-linking of mNb6 with the Spike RBD over the original FSY. In a recent communication,^47^ Han *et al*. exploited the FSY and PERx mechanism developed by us^19,24^ and incorporated FSY into the Spike-specific minibinders *de novo* designed by the Baker group.^8^ The resultant FSY-minibinder required 2 h incubation to crosslink with the Spike RBD *in vitro*, and 2 h incubation with virus to show a moderate 6-fold increase in potency in neutralizing a single SARS-CoV-2 variant. It is unclear if the potency increase by the FSY-minibinder is general, as no neutralization of WT SARS-CoV-2 or other variants was demonstrated. The longer the incubation time required for the drug, the more likely the virus would gain access to human cells for infection. In stark contrast, our new generation of FFY-enabled covalent nanobody mNb6(108FFY) exhibited robust cross-linking with the Spike RBD within 10 min. We incubated mNb6(108FFY) with SARS-CoV-2 for a shorter 1 h, and achieved a drastic 41-fold increase in potency in neutralizing SARS-CoV-2. In addition, we showed that mNb6(108FFY) also increased the potency in neutralizing the WT, Alpha, and Delta variant of SARS-CoV-2 pseudovirus by 36-, 23-, and 39-fold, respectively, demonstrating that the potency increase by the covalent nanobody was robust and general. Therefore, fast reaction kinetics of FFY is critical in enhancing the potency of covalent protein drugs.

We also uncovered that potency increase by covalent protein drug in neutralizing SARS-CoV-2 was dependent on the Spike RBD state that the protein drug binds. A puzzling observation was that a striking increase in antagonizing the interaction of Spike RBD with ACE2 *in vitro* by a covalent binder does not necessarily translate into more potent neutralization of viral infection. Compared with the WT nanobody SR4, the covalent nanobody SR4(57FSY) inhibited the Spike RBD binding to cell surface ACE2 with an enhanced potency of 125-fold, but it showed no enhancing effect in neutralizing SARS-CoV-2. We discovered that the underlying reason was that SR4 binds the Spike RBD in the latter’s active up state.^23^ In contrast, mNb6 binds the Spike RBD in the latter’s inactive down state,^7^ and the covalent nanobody mNb6(108FFY) then drastically increased the potency in neutralizing the WT, Alpha, and Delta variant of SARS-CoV-2. Therefore, to achieve potent neutralization of virus, the covalent binder needs to directly block the viral infection process aside from binding with the viral protein. This mechanistic insight will be valuable in guiding the development of covalent inhibitors for other viral infections.

Covalent nanobodies may lead to new protein drugs that can be readily produced in large scale via bacterial expression, easy to store and distribute due to nanobody’s high stability, and aerosolized for self-administered inhalation to nasal and lung epithelia.^7^ Further development may afford a medication for COVID-19 patients to prevent significant morbidities and death, and provide a potential prophylactic to give passive immunity to clinical providers at the front line. In addition, the PERx-capable ACE2 drugs can serve as a therapeutic stockpile for future outbreaks of SARS-CoV, SARS-CoV-2, and any new coronavirus or its variants that use the ACE2 receptor for entry. Moreover, through irreversible binding covalent protein drugs can potentially achieve complete viral inhibition and mechanistically prevent viral resistance. Lastly, the principle of generating a covalent binder or a covalent soluble receptor inhibitor via PERx can be generally applied for developing covalent protein drugs to effectively treat various other infectious diseases such as influenza, hepatitis, AIDS, anthrax, and so on.

## EXPERIMENTAL PROCEDURES

### Nanobody expression and purification

Plasmid pBAD-H11D4, pBAD-MR17K99Y, or pBAD-SR was transformed into *E. coli* BL21(DE3) for wildtype protein expression. Plasmid pBAD-H11D4 (27TAG), pBAD-H11D4 (30TAG), pBAD-H11D4 (100TAG), pBAD-H11D4 (112TAG), pBAD-H11D4 (115TAG), pBAD-H11D4 (116TAG), pBAD-MR17K99Y (99TAG), pBAD-MR17K99Y (101TAG), pBAD-SR4 (37TAG), pBAD-SR4(54TAG), or pBAD-SR4 (57TAG) was co-transformed respectively with plasmid pEVOL-FSYRS into *E. coli* BL21(DE3) for FSY incorporated protein expression. Transformed *E*.*coli* cells were grown at 37 °C, 220 rpm to an OD 0.5, after which 0.2 % L-arabinose was added. For FSY incorporation, 1 mM FSY was added to the growth medium. The expression was carried out at 18 °C, 220 rpm for 18 h. Cells were harvested at 8000 g, 4 °C for 30 min. The cell pellet was suspended with lysis buffer (20 mM Tris-HCl, 20 mM imidazole, 200 mM NaCl, pH 7.5) supplemented with EDTA free protease inhibitor cocktail and 1 μg/mL DNase. The cells were lysed by sonification, after which the cell lysis solution was centrifuged at 25,000 g at 4 °C for 40 min. The supernatant was collected and purified with 500 μL Ni-NTA resin affinity resin. The resin was washed and eluted with elution buffer (20 mM Tris-HCl, 300 mM imidazole, 200 mM NaCI, pH 7.5). The eluted protein was concentrated and exchanged with buffer (20 mM Tris-HCl, 200 mM NaCI, pH 7.5). To yield pure protein, TALON® metal affinity resin was further applied. The purification procedure was same as described above. The eluted proteins were analyzed by running 12% Tris-glycine SDS−PAGE gel.

### Cross-linking study of nanobody with Spike RBD

The Spike RBD was incubated with wildtype nanobody or nanobody mutants at indicated concentrations in PBS (pH 7.4) at 37 °C for 12 h, after which 5 μL sample was mixed with 10 μL Laemmli loading dye supplied with 100 mM DTT. The mixture was boiled at 95 °C for 10 min and subjected to Western blot analysis. The bands were detected using an antibody against mouse Fc appended at the C-terminus of the Spike RBD.

### Cellular surface ACE2 binding competition assay

Various concentrations of SR4 or SR4(57FSY) (final conc. 2, 1, 0.2, 0.05, 0.01, and 0.002 μM) were mixed with Spike RBD-mFc (final conc. 10 nM) in a final volume of 10 μL HBSS for overnight incubation at 37 °C. The mixture was diluted 50 times with HBSS and incubated at 37 °C for 1 h, 100 μL of which was subsequently added to 1.5 × 10^5^ ACE2-expressing 293T cells and incubated at 37 °C for 1 h. The cells were then washed twice with HBSS and labeled with an FITC-conjugated mFc-specific antibody at RT for 1 h, after which the cells were washed and analyzed with flow cytometry.

### Cross-linking kinetic study of Spike RBD with mNb6(108FFY) or mNb6(108FSY)

The SARS-CoV-2 WT or variant Spike RBD (Hisx6 tagged, 0.5 μM) was incubated with 5 μM mNb6(108FSY) or mNb6(108FFY) in PBS (pH 7.4) at 37 °C. At different time points, 5 μL reaction mixture was extracted and mixed with 5 μL Laemmli loading buffer. The mixture was heated to 95 °C for 10 min and the protein cross-linking was examined by Western blot. The protein band was detected with HRP-conjugated anti-Hisx6 antibody (Proteintech, #HRP-66005). The Spike RBD band intensity in the Western blot was quantified with Bio-rad imaging software. The linear plot of natural logarithm (ln) of the Spike RBD band intensity versus time (h) gives *k*_obs_.

### Binding constant (*K*_D_) measurement between Spike RBD and nanobody

Binding constant (*K*_D_) between Spike protein RBD and nanobody was measured with biolayer interferometry (BLI) using Octet Red384 systems (ForteBio). Biotinylated Spike RBD was firstly loaded to streptavidin (SA) sensor (ForteBio #18-5019) by incubating SA sensor in 100 nM biotinylated Spike RBD in Kinetic Buffer (0.005 % (v/v) Tween 20 and 0.1 % BSA in PBS, pH = 7.4) at 25 °C. The sensor was equilibrated (baseline step) in Kinetic Buffer for 120 s, after which the sensor was incubated with varying concentrations of nanobody (association step) for 50 s, followed with dissociation step in Kinetic Buffer for 450 s. Data was fitted for a 1:1 stoichiometry and *K*_D_ was calculated using the built in software.

### SARS-CoV-2 pseudovirus assay for nanobody neutralization

The SARS-CoV-2 GFP Reporter Virus Particles (RVPs) are SARS-CoV-2 pseudotyped lentivirus and were purchased from Integral Molecular. Catalog numbers for strains are the following: wild-type strain: RVP-701; Alpha: RVP-706; Beta: RVP-714; Delta: RVP-763. One day before transduction, 4 × 10^4^ 293T-ACE2 cells were plated in each well of a 48-well plate. Serially diluted nanobodies were incubated with pseudovirus in DMEM at 37 °C for 1 h. The mixture was subsequently transferred to each well of the 48-well plate. The cells were cultured at 37 °C for additional 48 h, after which the cells were harvested for flow cytometric analysis to measure the proportion of GFP positive cells.

### Authentic SARS-CoV-2 neutralization assay

SARS-CoV-2 Nanoluciferase (USA/WA1-2020) (SARS-CoV2 nLuc) was a kind gift from Dr. Pei-Yong Shi. The virus stocks were prepared in Vero E6 (ATCC) and titers were determined by plaque assays on Vero-E6 cells. Neutralizing assays were performed in 293T-ACE2 cells (15,000 cells per well) in a white opaque 96-well plates. Input virus (multiplicity of infection 0.01) was mixed with media containing nanobodies at indicated final concentrations and incubated for 1 h at 37 °C. The virus-nanobody mixtures were then added to cells for 2 h to allow virus adsorption, and washed. At 72 h post-infection, NanoLuc Luciferase substrates (Promega) was added to each well and luciferase signals were measured using a GloMax microplate reader (Promega). The relative luciferase signals were normalized to no-nanobody control. Virus propagation and experiments were performed in the BSL3 facility of the Gladstone Institutes with approved protocols.

### ACE2(FSY) expression and purification

One day before transfection, 2 × 10^6^ Expi293F™ cells were seeded with pre-warmed Freestyle 293 media. The cells were transfected with 25 μg pcDNA-ACE2-34TAG and 25 μg pMP-FSYRS plasmids according to manufacturer’s protocol. The cells were incubated for 30 min and then 2 mM FSY was added into the culture media dropwise. The supernatant was collected after 4 days post-transfection. Imidazole (20 mM) and pre-equilibrated Ni-NTA resin were added into the supernatant, followed by incubation 4 °C on the rotator for 1 h. The resin was washed and eluted with elution buffer (20 mM Tris-HCl, 300 mM imidazole, 200 mM NaCI, pH 7.5). The eluted protein was concentrated and exchanged with PBS, pH 7.5.

## ACKNOWLEDGEMENTS

L.W. acknowledges the support of UCSF Catalyst Award (7030487) and the NIH (R01GM118384). M.O. thanks the Rodenberry foundation, NIH grant R37AI083139, and the Gladstone Institutes for their support.

