## Supplementary Information for "Accelerating PERx Reaction Enables Covalent Nanobodies for Potent Neutralization of SARS-Cov-2 and Variants"

**Key reagents**

The SARS-CoV-2 GFP Reporter Virus Particles (RVPs, RVP-701G), Alpha variant RVPs RVP-706G), Beta variant RVPs (RVP-714G) and Delta variant RVPs (RVP-763G) were purchased form Integral Molecular, Inc. SARS-CoV-2 Spike RBD-mFc Recombinant Protein (40592-V05H) was purchased from SinoBiological company. Biotinylated SARS-CoV-2 Spike RBD (SPD-C82E9), biotinylated SARS-CoV-2 Spike RBD (N501Y) (SPD-C82E6), biotinylated SARS-CoV-2 Spike RBD (K417N, E484K, N501Y) (SPD-C82E5), biotinylated SARS-CoV-2 Spike RBD (L452R, E484K) (SPD-C82Ed) were purchased from ACROBiosystems. SARS SARS-CoV-2 Spike RBD (SPD-C52H3), SARS-CoV-2 Spike RBD (N501Y) (SPD-C52Hn), SARS-CoV-2 Spike RBD (K417N, E484K, N501Y) (SPD-C52Hp), SARS-CoV-2 Spike RBD (L452R, E484K) (SPD-C52Hh) were purchased from ACROBiosystems.

**Cell culture**

The 293T-hsACE2 stable cell line was purchased from Integral Molecular, Inc (C-HA102). The cell line was maintained in DMEM with 10 % FBS, 1 x penicillin-streptomycin and 0.5 µg/ml puromycin.

**Primers**

| | |
|---|---|
| H11D4-F | TAAGAAGGAGATATACATATGAAATATCTGCTGCCAACCG |
| H11D4-R | GCCAAAACAGCCAAGCTTTTAATGATGGTGATGGTGGTGTT |
| H11D4-R27TAG-F | GGTTAGCGGTTAGACCTTTAGCACC |
| H11D4-R27TAG-R | GCGCAACTCAGACGCAGA |
| H11D4-S30TAG-F | TCGCACCTTTTAGACCGCCGCGATGGGTTG |
| H11D4-S30TAG-R | CCGCTAACCGCGCAACTC |
| H11D4-E100TAG-F | CGCGCGCACCTAGAACGTTCGTA |
| H11D4-E100TAG-R | CAATAGTACACGGCGGTG |
| H11D4-W112TAG-F | TTACGCCACGTAGCCGTACGATT |
| H11D4-W112TAG-R | TCGCTCAGCAGACTACGAAC |

| | |
|---|---|
| H11D4-D115TAG-F | GTGGCCGTACTAGTACTGGGGTC |
| H11D4-D115TAG-R | GTGGCGTAATCGCTCAGC |
| H11D4-Y116TAG-F | GCCGTACGATTAGTGGGGTCAAG |
| H11D4-Y116TAG-R | CACGTGGCGTAATCGCTC |
| MR17K99Y-F | CTTTAAGAAGGAGATATACATATGAAATATCTGCTGCCAACCG |
| MR17K99Y-R | TCCGCCAAAACAGCCAAGCTTTTAATGGTGATGATGATGG |
| MR17K99Y-Y99TAG-F | CTGCAACGTGTAGGATGATGGCC |
| MR17K99Y-Y99TAG-R | TAGTACACGGCCGTATCC |
| MR17K99Y-D101TAG-F | CGTGTACGATTAGGGCCAGCTGG |
| MR17K99Y-D101TAG-R | TTGCAGTAGTACACGGCC |
| SR4-F | CTTTAAGAAGGAGATATACATATGAAATATCTGCTGCCAACCGC |
| SR4-R | ATCCGCCAAAACAGCCAAGCTTTTAATGATGATGGTGATGG |
| SR4-Y37TAG-F | CATGTGGTGGTAGCGCCAAGCCC |
| SR4-Y37TAG-R | TTCCAGCTGTACACTGGAAAGCC |
| SR4-H54TAG-F | GATCGAAAGCTAGGGCGATAGCACCC |
| SR4-H54TAG-R | GCCGCAACCCATTCGCGT |
| SR4-S57TAG-F | CCACGGCGATTAGACCCGCTACGCG |
| SR4-S57TAG-R | CTTTCGATCGCCGCAACC |
| pcDNA-ACE2-Hind3-F | CTAGCGTTTAAACTTAAGCTTGCCACCATGTCAAGCTCTTCCTGGCTC |
| pcDNA-ACE2-His-BamHI-R | CACACTGGACTAGTGGATCCTTAGTGATGGTGATGATGATGGGAAACAGGGGGCTGG |
| ACE2-D30TAG-F | CCAAGACATTTTTGTAGAAGTTTAACCACG |
| ACE2-D30TAG-R | CTACAAAAATGTCTTGGCCTGTTCCTC |
| ACE2-D38TAG-F | GTTTAACCACGAAGCCGAATAGCTGTTCTATCAAAG |
| ACE2-D38TAG-R | CTATTCGGCTTCGTGGTTAAACTTG |

| | |
|---|---|
| ACE2-E37TAG-F | CAAGTTTAACCACGAAGCCTAGGACCTGTTCTATCAAAG |
| ACE2-E37TAG-R | CTAGGCTTCGTGGTTAAACTTGTC |
| ACE2-H34TAG-F | CATTTTTGGACAATTTAACTAGGAAGCCGAAGACCTG |
| ACE2-H34TAG-R | GGCTTCCTAGTTAAACTTGTCCAAAAATG |
| ACE2-Q42TAG-F | CGAAGACCTGTTCTATTAGAGTTCACTTGCTTC |
| ACE2-Q42TAG-R | CTCTAATAGAACAGGTCTTCGGCTTCGTG |
| SDM-ACE2-83TAG-F | TGCCCAAATGTAGCCACTACAAG |
| SDM-ACE2-83TAG-R | AGTGTGGACTGTTCCTTTAAAAAG |
| mNb6-F | TTTAACTTTAAGAAGGAGATATACATATGAAATATCTGCTGCCAACCGCGGC |
| mNb6-R | ATCCGCCAAAACAGCCAAGCTTTTAATGATGATGGTGATGGTGGCTGCTCA |
| mNb6-G26TAG-F | CGGCGAGCTAGTATATTTTTGGCCGCAACGCG |
| mNb6-G26TAG-R | AATATACTAGCTCGCCGCGCAGCTCAGGCGCA |
| mNb6-Y27TAG-F | GAGCGGCTAGATTTTTGGCCGCAACGCGATGG |
| mNb6-Y27TAG- | CAAAAATCTAGCCGCTCGCCGCGCAGCTCAGG |
| mNb6-I28TAG-F | CGGCTATTAGTTTGGCCGCAACGCGATGGGCT |
| mNb6-I28TAG-R | GGCCAAACTAATAGCCGCTCGCCGCGCAGCTC |
| mNb6-F29TAG-F | CTATATTTAGGGCCGCAACGCGATGGGCTGGT |
| mNb6-F29TAG- | TGCGGCCCTAAATATAGCCGCTCGCCGCGCAG |
| mNb6-G30TAG-F | TTTTTTAGCGCAACGCGATGGGCTGGTATCGC |
| mNb6-G30TAG-R | CGCGTTGCGCTAAAAAATATAGCCGCTCGCCG |
| mNb6-R31TAG-F | TTTTGGCTAGAACGCGATGGGCTGGTATCGCC |
| mNb6-R31TAG-R | TCGCGTTCTAGCCAAAAATATAGCCGCTCGCC |
| mNb6-N32TAG-F | ATTTTTGGCCGCTAGGCGATGGGCTGGTATCGC |

| | |
|---|---|
| mNb6-N32TAG-R | GCCTAGCGGCCAAAAATATAGCCGCTCGCCGC |
| mNb6-A33TAG-F | CAACTAGATGGGCTGGTATCGCCAAGCGCCGG |
| mNb6-A33TAG-R | ACCAGCCCATCTAGTTGCGGCCAAAAATATAGCC |
| mNb6-M34TAG-F | AACGCGTAGGGCTGGTATCGCCAAGCGCCGGG |
| mNb6-M34TAG-R | TACCAGCCCTACGCGTTGCGGCCAAAAATATA |
| mNb6-G35TAG-F | CGCGATGTAGTGGTATCGCCAAGCGCCGGGCA |
| mNb6-G35TAG-R | GATACCACTACATCGCGTTGCGGCCAAAAATA |
| mNb6-G50TAG-F | ACTGGTGGCGTAGATTACCCGCCGC |
| mNb6-G50TAG-R | TCGCGTTCTTTGCCCGGC |
| mNb6-I51TAG-F | GGTGGCGGGCTAGACCCGCCGCG |
| mNb6-I51TAG-R | AGTTCGCGTTCTTTGCCCGG |
| mNb6-T52TAG-F | GGCGGGCATTTAGCGCCGCGGCA |
| mNb6-T52TAG-R | ACCAGTTCGCGTTCTTTGCC |
| mNb6-R53TAG-F | GGGCATTACCTAGCGCGGCAGCATTAC |
| mNb6-R53TAG-R | GCCACCAGTTCGCGTTCT |
| mNb6-R54TAG-F | CATTACCCGCTAGGGCAGCATTACCTATTATGCGGATAG |
| mNb6-R54TAG-R | CCCGCCACCAGTTCGCGT |
| mNb6-G55TAG-F | TACCCGCCGCTAGAGCATTACCTATTATGC |
| mNb6-G55TAG-R | ATGCCCGCCACCAGTTCG |
| mNb6-S56TAG-F | CCGCCGCGGCTAGATTACCTATTATGCGG |
| mNb6-S56TAG- | GTAATGCCCGCCACCAGT |
| mNb6-I57TAG-F | CCGCGGCAGCTAGACCTATTATGCGGATAG |
| mNb6-I57TAG-R | CGGGTAATGCCCGCCACC |

| | |
|---|---|
| mNb6-T58TAG-F | CGGCAGCATTTAGTATTATGCGGATAGCGTGAAAGGCCGC |
| mNb6-T58TAG-R | CGGCGGGTAATGCCCGCC |
| mNb6-Y59TAG-F | CAGCATTACCTAGTATGCGGATAGCGTGAAAGG |
| mNb6-Y59TAG-R | CCGCGGCGGGTAATGCCC |
| mNb6-D99TAG-F | ATTATTGCGCGGCGTAGCCGGCGAGCCCGGCGTAT |
| mNb6-D99TAG-R | CTACGCCGCGCAATAATACACCGCGGTATCTT |
| mNb6-P100TAG-F | ATTGCGCGGCGGATTAGGCGAGCCCGGCGTATGGC |
| mNb6-P100TAG-R | CTAATCCGCCGCGCAATAATACACCGCGGTAT |
| mNb6-A101TAG-F | CGTAGAGCCCGGCGTATGGCGATTATTGGGGC |
| mNb6-A101TAG-R | ATACGCCGGGCTCTACGGATCCGCCGCGCAATA |
| mNb6-S102TAG-F | TAGCCGGCGTATGGCGATTATTGGGGCCAAGG |
| mNb6-S102TAG-R | TCGCCATACGCCGGCTACGCCGGATCCGCCGCGCA |
| mNb6-P103TAG-F | CTAGGCGTATGGCGATTATTGGGGCCAAGGCA |
| mNb6-P103TAG-R | AATCGCCATACGCCTAGCTCGCCGGATCCGCCGC |
| mNb6-A104TAG-F | CCCGTAGTATGGCGATTATTGGGGCCAAGGCA |
| mNb6-A104TAG-R | AATCGCCATACTACGGGCTCGCCGGATCCGCC |
| mNb6-Y105TAG-F | GGCGTAGGGCGATTATTGGGGCCAAGGCACCC |
| mNb6-Y105TAG-R | AATAATCGCCCTACGCCGGGCTCGCCGGATCC |
| mNb6-G106TAG-F | CCGGCGTATTAGGATTATTGGGGCCAAGGCAC |
| mNb6-G106TAG-R | TAATCCTAATACGCCGGGCTCGCCGGATCCGC |
| mNb6-D107TAG-F | GTATGGCTAGTATTGGGGCCAAGGCACCCAAG |
| mNb6-D107TAG-R | CCCAATACTAGCCATACGCCGGGCTCGCCGGA |
| mNb6-Y108TAG-F | TATGGCGATTAGTGGGGCCAAGGCACCCAAGT |

| | |
|---|---|
| mNb6-<br>Y108TAG-R | CCCCACTAATCGCCATACGCCGGGCTCGCCGG |

**Molecular cloning**

H11-D4, MR17-K99Y and SR4 and mNb6 fragment genes were ordered from Genewiz. Human ACE2 gene was ordered from Addgene #1786.

pBAD-H11D4: primers H11D4-F and H11D4-R were used to amplify H11D4 fragments. pBAD vector and H11D4 fragment were joined with recombination cloning to create the pBAD-H11D4. Using pBAD-H11D4 as template, primers H11D4-R27TAG-F and H11D4-R27TAG-R were used to generate pBAD-H11D4 (27TAG); primers H11D4-S30TAG-F and H11D4-S30TAG-R were used to generate pBAD-H11D4 (30TAG); primers H11D4-E100TAG-F and H11D4-E100TAG-R were used to generate pBAD-H11D4 (100TAG); primers H11D4-W112TAG-F and H11D4-W112TAG-R were used to generate pBAD-H11D4 (112TAG); primers H11D4-D115TAG-F and H11D4-D115TAG-R were used to generate pBAD-H11D4 (115TAG); primers H11D4-Y116TAG-F and H11D4-Y116TAG-R were used to generate pBAD-H11D4 (116TAG). Amino acid sequence of cpsGFP with Tyr66 highlighted in red is shown below.

pBAD-MR17K99Y: MR17K99Y-F and MR17K99Y-R were used to amplify MR17K99Y fragments. pBAD vector and MR17K99Y fragment were joined with recombination cloning to create the pBAD-MR17K99Y. Using pBAD-MR17K99Y as template, primers MR17K99Y-Y99TAG-F and MR17K99Y-Y99TAG-R were used to generate pBAD-MR17K99Y (99TAG); primers MR17K99Y-D101TAG-F and MR17K99Y-D101TAG-R were used to generate pBAD-MR17K99Y (101TAG).

pBAD-SR4: SR4-F and SR4-R were used to amplify SR4 fragments. pBAD vector and SR4 fragment were joined with recombination cloning to create the pBAD-SR4. Using pBAD-SR4 as template, primers SR4-Y37TAG-F and SR4-Y37TAG-R were used to generate pBAD-SR4 (37TAG); primers SR4-H54TAG-F and SR4-H54TAG-R were used to generate pBAD-SR4 (54TAG); primers SR4-S57TAG-F and SR4-S57TAG-R were used to generate pBAD-SR4 (57TAG).

pcDNA3.1-ACE2: pcDNA-ACE2-Hind3-F and pcDNA-ACE2-His-BamHI-R were used to amplify the ACE2 gene. pcDNA3.1 vector and ACE2 fragment were joined with recombination cloning to create the pcDNA3.1-ACE2. Using pcDNA3.1-ACE2 as template, primers with TAG were used to generate its TAG mutants.

pBAD-mNb6: mNb6-F and mNb6-R were used to amplify mNb6 fragments. pBAD vector and mNb6 fragment were joined with recombination cloning to create the pBAD-mNb6. Using pBAD-mNb6 as template, primers with TAG were used to generate its TAG mutants.

Wildtype H11D4 nanobody gene seuqnence:

ATGAAATATCTGCTGCCAACCGCGGCCGCGGGTCTGCTGCTGCTGGCGGCCCAACCAGCGATGG
CGCAAGTGCAGCTGGTTGAGAGCGGCGGTGGTCTGATGCAAGCGGGTGGTAGTCTGCGTCTGAG
TTGCGCGGTTAGCGGTCGCACCTTTAGCACCGCCGCGATGGGTTGGTTTCGCCAAGCGCCGGGC
AAAGAACGCGAATTTGTTGCGGCCATCCGTTGGAGCGGTGGTAGTGCGTACTACGCGGATAGCG
TTAAAGGCCGCTTCACCATCAGCCGCGATAAGGCGAAGAACACCGTGTATCTGCAGATGAACAG
TCTGAAGTACGAGGACACCGCCGTGTACTATTGCGCGCACCGAAAACGTTCGTAGTCTGCTG
AGCGATTACGCCACGTGGCCGTACGATTACTGGGGTCAAGGCACCCAAGTTACCGTGAGCAGCA
AACACCACCATCACCATCAT

Wildtype MR17K99Y nanobody gene seuqnence:

ATGAAATATCTGCTGCCAACGGCCGCGGCGGGTCTGCTGCTGCTGGCGGCGCAACCAGCGATGG
CCCAAGTTCAGCTGGTTGAAAGCGGTGGCGGTCTGGTTCAAGCCGGTGGTAGTCTGCGTCTGAG
CTGCGCCGCCAGTGGCTTTCCGGTGGAAGTTTGGCGCATGGAATGGTACCGCCAAGCCCCGGGC
AAAGAACGCGAAGGCGTTGCCGCCATCGAAAGCTACGGTCATGGCACCCGCTACGCCGATAGCG
TTAAAGGCCGCTTCACCATCAGCCGCGACAACGCGAAGAACACCGTGTATCTGCAGATGAACAG
TCTGAAACCGGAGGATACGGCCGTGTACTACTGCAACGTGTACGATGATGGCCAGCTGGCGTAC
CATTACGATTACTGGGGCCAAGGCACCCAAGTTACCGTTAGTGCGGGTCGCGCGGGCGAACAGA
AGCTGATCAGCGAAGAGGATCTGAATAGCGCCGTGGATCACCATCATCATCACCAT

Wildtype SR4 nanobody gene seuqnence:

ATGAAATATCTGCTGCCAACCGCCGCGGCGGGTCTGCTGCTGCTGGCGGCGCAACCAGCGATGG
CCCAAGTTCAGCTGGTTGAAAGCGGTGGTGGTCTGGTTCAAGCCGGTGGTAGTCTGCGTCTGAG
CTGCGCGGCGAGTGGCTTTCCAGTGTACAGCTGGAACATGTGGTGGTACCGCCAAGCCCCGGGT
AAAGAACGCGAATGGGTTGCGGCGATCGAAAGCCACGGCGATAGCACCCGCTACGCGGATAGCG
TTAAAGGCCGCTTCACCATCAGCCGCGACAACGCCAAGAACACCGTGTATCTGCAGATGAACAG
TCTGAAACCGGAAGATACCGCGGTGTACTACTGCTACGTGTGGGTTGGCCACACCTACTACGGT

CAAGGCACCCAAGTTACCGTTAGCGCGGGTCGTGCGGGCGAACAGAAGCTGATCAGCGAGGAAG
ATCTGAACAGCGCCGTGGATCACCATCACCATCATCAT

Wildtype ACE2 gene seuqnence:

ATGTCAAGCTCTTCCTGGCTCCTTCTCAGCCTTGTTGCTGTAACTGCTGCTCAGTCCACCATTG
AGGAACAGGCCAAGACATTTTTGGACAAGTTTAACCACGAAGCCGAAGACCTGTTCTATCAAAG
TTCACTTGCTTCTTGGAATTATAACACCAATATTACTGAAGAGAATGTCCAAAACATGAATAAT
GCTGGGGACAAATGGTCTGCCTTTTTAAAGGAACAGTCCACACTTGCCCAAATGTATCCACTAC
AAGAAATTCAGAATCTCACAGTCAAGCTTCAGCTGCAGGCTCTTCAGCAAAATGGGTCTTCAGT
GCTCTCAGAAGACAAGAGCAAACGGTTGAACACAATTCTAAATACAATGAGCACCATCTACAGT
ACTGGAAAAGTTTGTAACCCAGATAATCCACAAGAATGCTTATTACTTGAACCAGGTTTGAATG
AAATAATGGCAAACAGTTTAGACTACAATGAGAGGCTCTGGGCTTGGGAAAGCTGGAGATCTGA
GGTCGGCAAGCAGCTGAGGCCATTATATGAAGAGTATGTGGTCTTGAAAAATGAGATGGCAAGA
GCAAATCATTATGAGGACTATGGGGATTATTGGAGAGGAGACTATGAAGTAAATGGGGTAGATG
GCTATGACTACAGCCGCGGCCAGTTGATTGAAGATGTGGAACATACCTTTGAAGAGATTAAACC
ATTATATGAACATCTTCATGCCTATGTGAGGGCAAAGTTGATGAATGCCTATCCTTCCTATATC
AGTCCAATTGGATGCCTCCCTGCTCATTTGCTTGGTGATATGTGGGGTAGATTTTGGACAAATC
TGTACTCTTTGACAGTTCCCTTTGGACAGAAACCAAACATAGATGTTACTGATGCAATGGTGGA
CCAGGCCTGGGATGCACAGAGAATATTCAAGGAGGCCGAGAAGTTCTTTGTATCTGTTGGTCTT
CCTAATATGACTCAAGGATTCTGGGAAAATTCCATGCTAACGGACCCAGGAAATGTTCAGAAAG
CAGTCTGCCATCCCACAGCTTGGGACCTGGGGAAGGGCGACTTCAGGATCCTTATGTGCACAAA
GGTGACAATGGACGACTTCCTGACAGCTCATCATGAGATGGGGCATATCCAGTATGATATGGCA
TATGCTGCACAACCTTTTCTGCTAAGAAATGGAGCTAATGAAGGATTCCATGAAGCTGTTGGGG
AAATCATGTCACTTTCTGCAGCCACACCTAAGCATTTAAAATCCATTGGTCTTCTGTCACCCGA
TTTTCAAGAAGACAATGAAACAGAAATAAACTTCCTGCTCAAACAAGCACTCACGATTGTTGGG
ACTCTGCCATTTACTTACATGTTAGAGAAGTGGAGGTGGATGGTCTTTAAAGGGGAAATTCCCA
AAGACCAGTGGATGAAAAAGTGGTGGGAGATGAAGCGAGAGATAGTTGGGGTGGTGGAACCTGT
GCCCCATGATGAAACATACTGTGACCCCGCATCTCTGTTCCATGTTTCTAATGATTACTCATTC
ATTCGATATTACACAAGGACCCTTTACCAATTCCAGTTTCAAGAAGCACTTTGTCAAGCAGCTA
AACATGAAGGCCCTCTGCACAAATGTGACATCTCAAACTCTACAGAAGCTGGACAGAAACTGTT
CAATATGCTGAGGCTTGGAAAATCAGAACCCTGGACCCTAGCATTGGAAAATGTTGTAGGAGCA
AAGAACATGAATGTAAGGCCACTGCTCAACTACTTTGAGCCCTTATTTACCTGGCTGAAAGACC

AGAACAAGAATTCTTTTGTGGGATGGAGTACCGACTGGAGTCCATATGCAGACCAAAGCATCAA
AGTGAGGATAAGCCTAAAATCAGCTCTTGGAGATAAAGCATATGAATGGAACGACAATGAAATG
TACCTGTTCCGATCATCTGTTGCATATGCTATGAGGCAGTACTTTTTAAAAGTAAAAAATCAGA
TGATTCTTTTTGGGGAGGAGGATGTGCGAGTGGCTAATTTGAAACCAAGAATCTCCTTTAATTT
CTTTGTCACTGCACCTAAAAATGTGTCTGATATCATTCCTAGAACTGAAGTTGAAAAGGCCATC
AGGATGTCCCGGAGCCGTATCAATGATGCTTTCCGTCTGAATGACAACAGCCTAGAGTTTCTGG
GGATACAGCCAACACTTGGACCTCCTAACCAGCCCCCTGTTTCCCATCATCATCACCATCAC

Wildtype mNb6 gene seuqnence:
ATGAAATATCTGCTGCCAACCGCGGCCGCGGGTCTGCTGCTGCTGGCGGCCCAACCAGCGATGG
CGCAAGTGCAGCTGGTGGAATCTGGCGGGGGCTTAGTGCAAGCGGGCGGCAGCCTGCGCCTGAG
CTGCGCGGCGAGCGGCTATATTTTTGGCCGCAACGCGATGGGCTGGTATCGCCAAGCGCCGGGC
AAAGAACGCGAACTGGTGGCGGGCATTACCCGCCGCGGCAGCATTACCTATTATGCGGATAGCG
TGAAAGGCCGCTTTACCATTAGCCGCGATAACGCGAAAAACACCGTGTATCTGCAGATGAACAG
CCTGAAACCGGAAGATACCGCGGTGTATTATTGCGCGGCGGATCCGGCGAGCCCGGCGTATGGC
GATTATTGGGGCCAAGGCACCCAAGTGACCGTGAGCAGCCACCATCACCATCATCAT

**Synthesis of FFY**

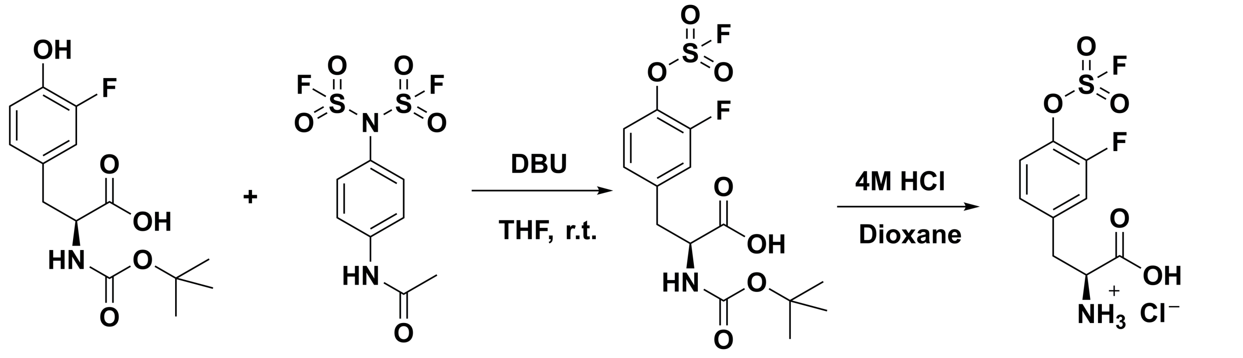


1 AISF 2 FFY

**Scheme 1. Synthesis of FFY.**

Synthesis of compound **2**. Compound **1** was converted to fluorosulfate using [4-(acetylamino)phenyl]imidodisulfuryl difluoride (AISF).[4] 1.0 g compound **1** (3.3 mmol) and AISF

(1.3 g, 4.0 mmol) was dissolved in 12 ml anhydrous THF. Then 1,8-Diazabicyclo[5.4.0]undec-7-ene (DBU, 1.1 g, 7.3 mmol) was added dropwise at room temperature (r.t.). The mixture was stirred at r.t. for 10 min. Then 100 ml EtOAc was added to dilute the reaction mixture and the organic phrase was washed sequentially by $H_2O$ (50 mL) and brine (50 mL). The organic phase was dried over anhydrous $Na_2SO_4$ and evaporated under reduced pressure to give the crude product, which was then purified by column chromatography (silica gel, DCM: MeOH=50:1) to give a white solid (0.8 g, 64 %).

Synthesis of **FFY**. Compound **2** (0.8g, 2.1 mmol) was stirred in 4 M HCl in dioxane (10 ml) at r.t. for 6 h. Then 10 ml diethyl ether was added to the reaction mixture, and a white precipitate was formed and collected by filtration. The white solid was further dried under reduced pressure to give **FFY** in HCl salt form (604 mg, 90 %). $^1$H NMR ($D_2O$): $\delta$ 7.63-7.59 (m, 1H), 7.39 (dd, $J$ = 10.8 Hz, $J$ = 2.0 Hz, 1H), 7.29-7.26 (m, 1H), 4.27-4.24 (m, 1H), 3.41-3.24 (m, 2H). $^{13}$C NMR ($D_2O$): $\delta$ 172.2, 153.8 (d, $J$ = 252 Hz, C-F), 138.4, 136.8 (d, $J$ = 13 Hz, C-F), 127.1 (d, $J$ = 3 Hz, C-F), 124.4, 119.4 (d, $J$ = 18 Hz, C-F), 54.8, 35.8; HRMS calcd for $C_9H_{10}F_2NO_5S$ $[M+H]^+$ 282.0242, found: 282.0253.

$^1$H NMR for FFY

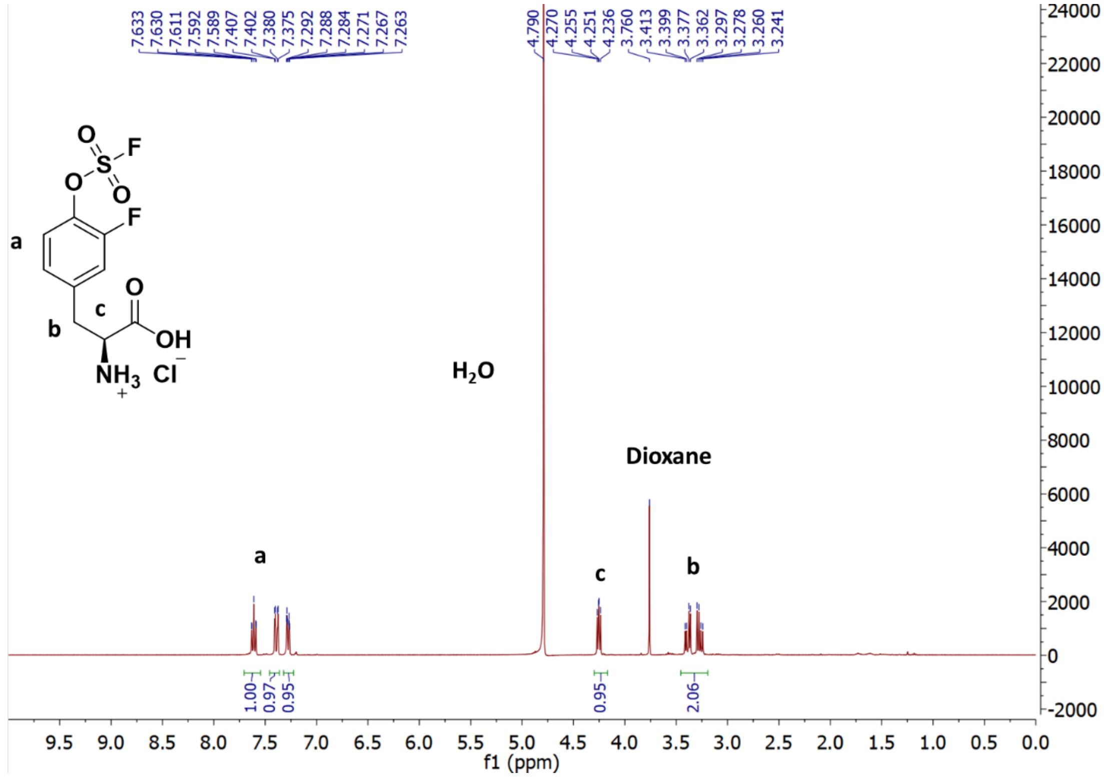

$^{13}C$ NMR for FFY

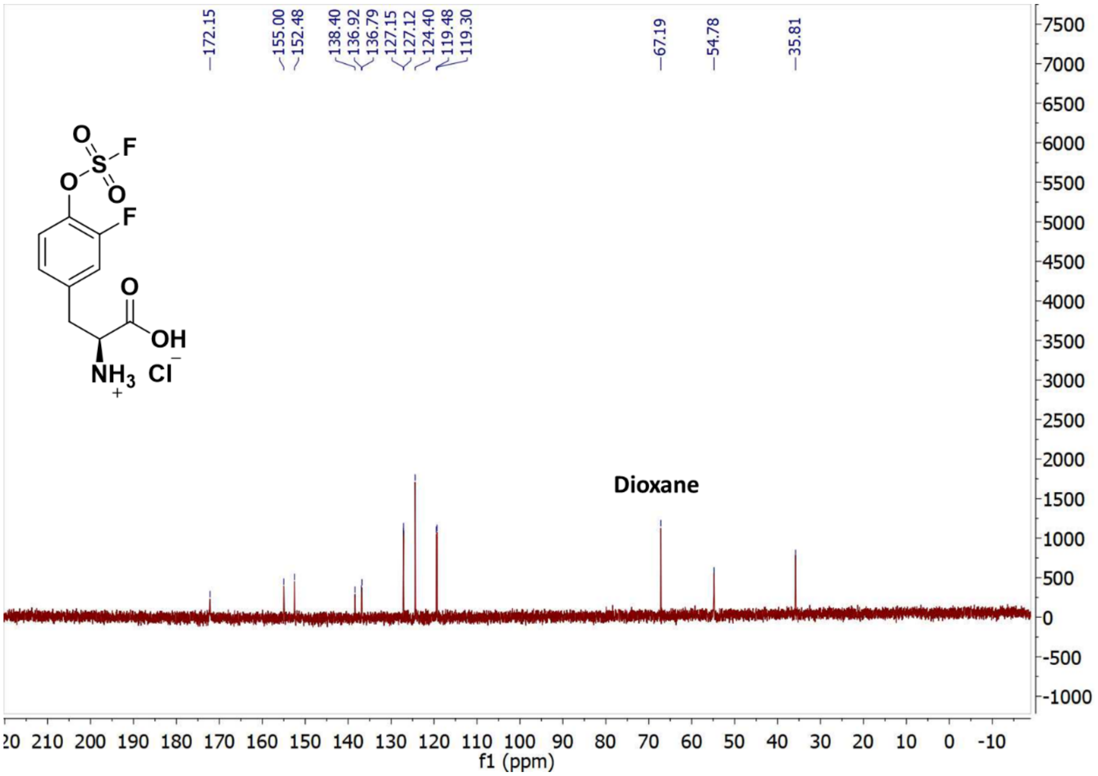

### Supplementary Figures

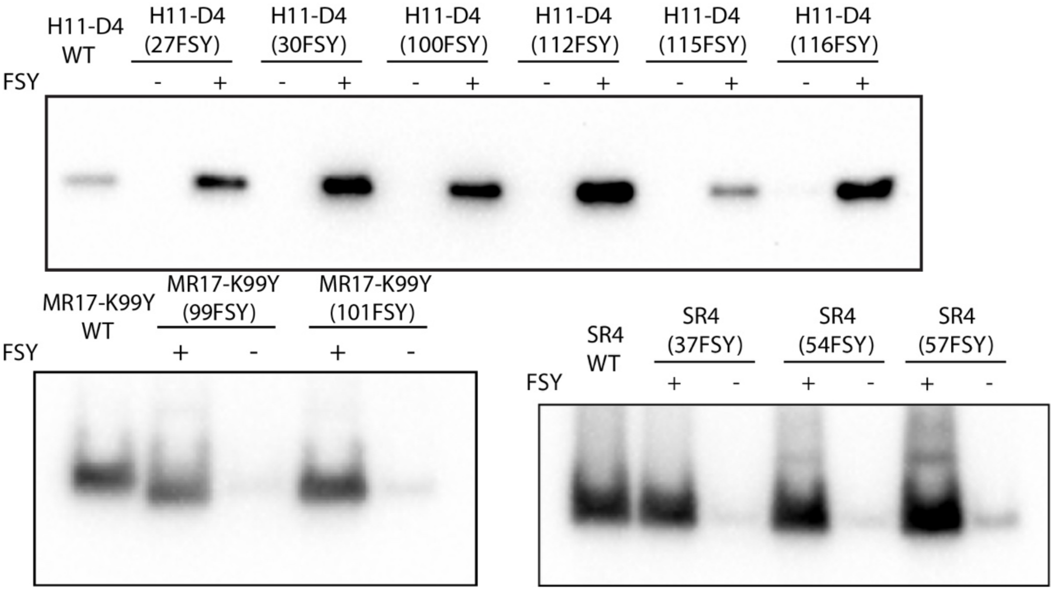


**Figure S1. Western blot analysis of the expression of nanobodies H11-D4, MR17-K99Y, SR4, and their FSY mutants.** For FSY mutants, full-length nanobodies were detected when FSY was added to the culture media, suggesting FSY incorporation. An antibody specific for the Hisx6 tag appended at the C-terminus of the nanobodies was used for detection.

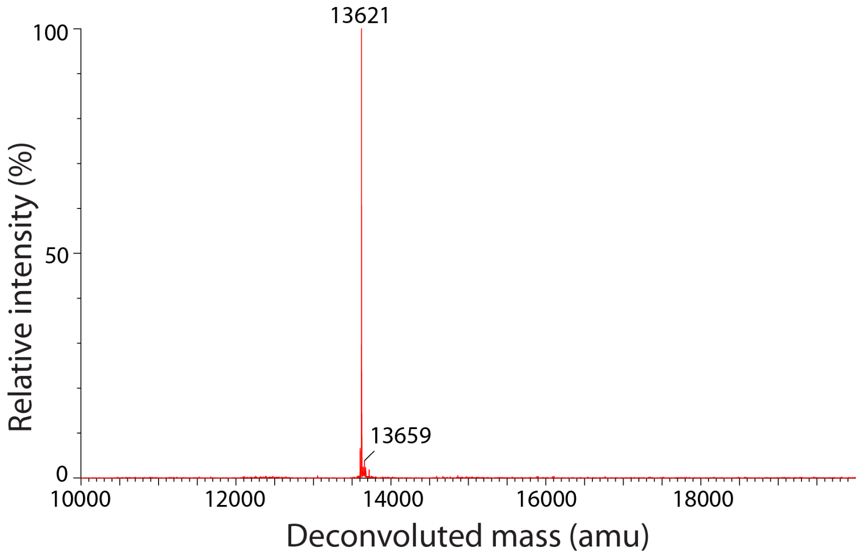


**Figure S2. Mass spectrometric analysis of the intact protein of mNb6(WT).** A major peak was observed at 13621 Da, corresponding to mNb6(WT) (expected at 13622.7 Da).

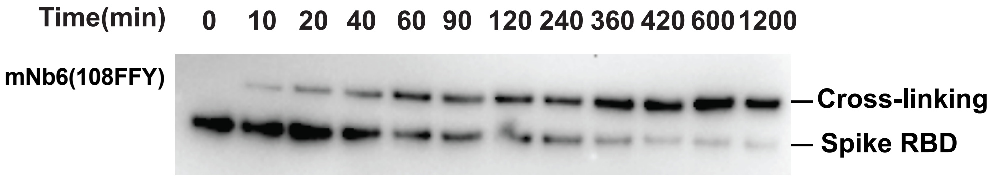


**Figure S3. Western blot analysis of mNb6(108FFY) (5 µM) cross-linking with the Spike RBD (0.5 µM) at indicated time points.** The bands were detected by an antibody against the mouse Fc tag appended on the Spike RBD. Cross-linking was robustly detected as early as 10 min.

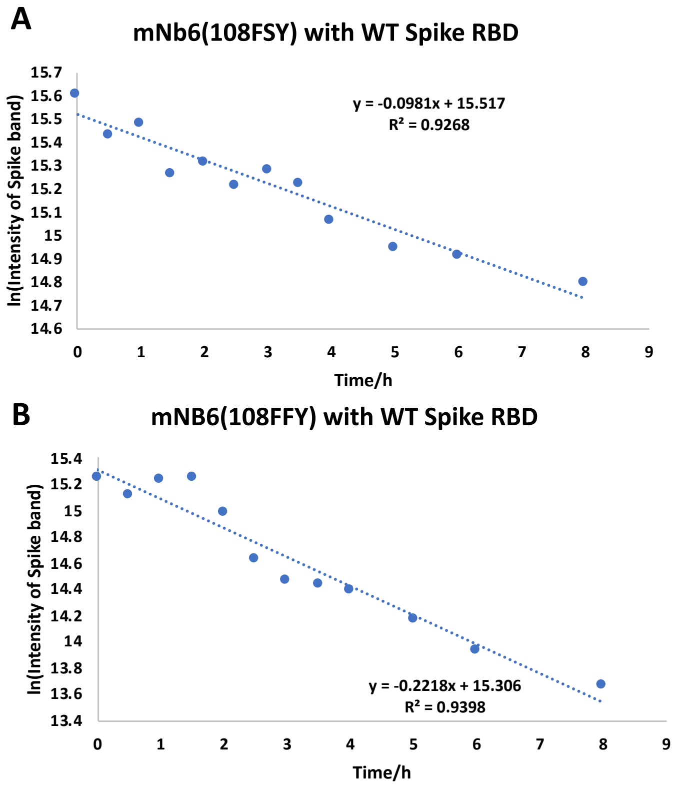


**Figure S4. Kinetics of mNB6(108FSY) or mNB6(108FFY) cross-linking with the WT Spike RBD *in vitro*.** $k_{obs}$ was calculated based on the time-dependent intensity decrease of the Spike RBD protein band. The band intensities in the Western blots were quantified with Bio-rad imaging software. The linear plot of natural logarithm (ln) of the Spike RBD band intensity versus time (h) gives $k_{obs}$. (**A**) $k_{obs}$ for mNb6(108FSY) with the WT Spike RBD was 0.0981 $h^{-1}$; (**B**) $k_{obs}$ for mNb6(108FFY) with the WT Spike RBD was 0.2218 $h^{-1}$. The experiments were independently repeated three times, and the data for one time are shown here.

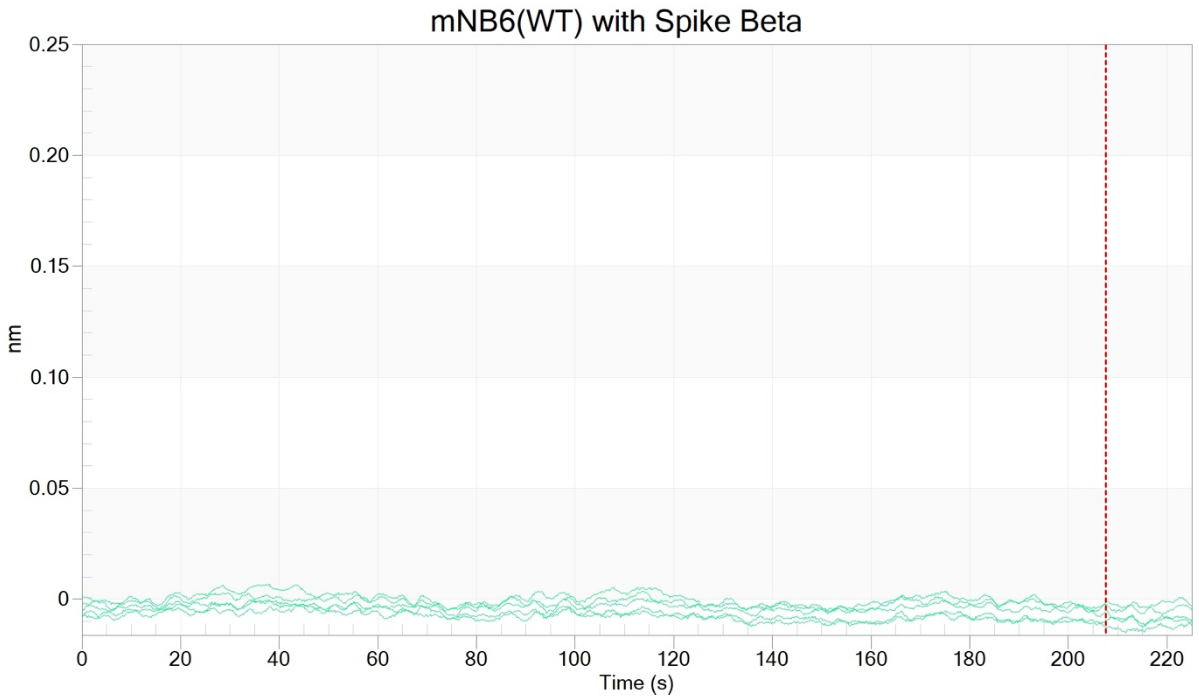


**Figure S5. BLI of mNb6(WT) bind to the Spike RBD of the Beta variant.** The concentrations for mNb6 were 0, 3.12, 6.25, 12.5, 25, and 50 nM. No significant binding was observed between mNb6(WT) and the Beta Spike RBD at these concentrations. The red vertical line indicates the beginning of dissociation step.

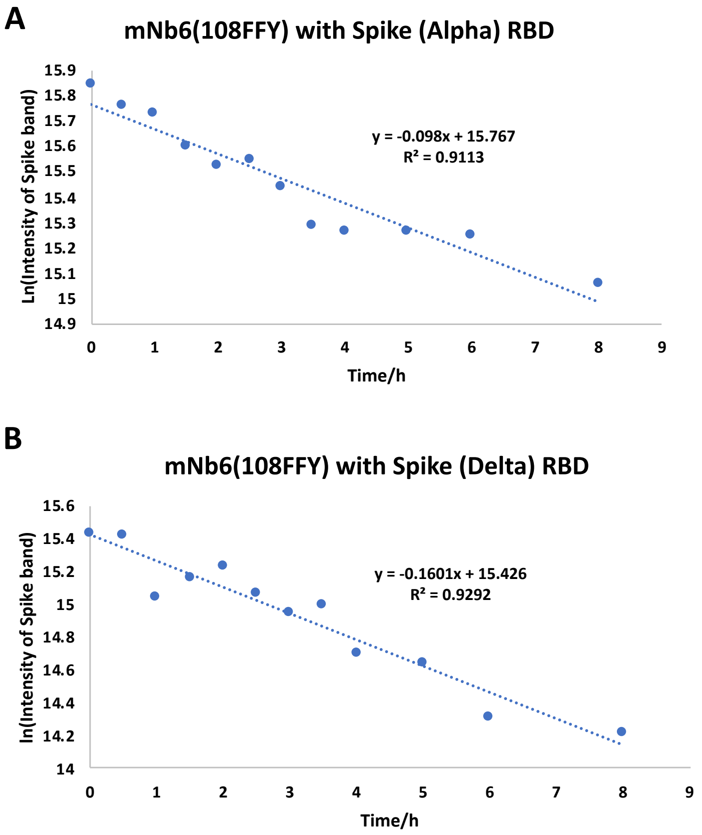


**Figure S6. Kinetics of mNB6(108FFY) cross-linking with the Spike RBD of SARS-CoV-2 variants *in vitro*.** $k_{obs}$ was calculated based on the time-dependent intensity decrease of the Spike RBD protein band. The band intensities in the Western blots were quantified with Bio-rad imaging software. The linear plot of natural logarithm (ln) of the Spike RBD band intensity versus time (h) gives $k_{obs}$. (**A**) $k_{obs}$ for mNb6(108FFY) with the Alpha Spike RBD was 0.098 $h^{-1}$; (**B**) $k_{obs}$ for mNb6(108FFY) with the Delta Spike RBD was 0.1601 $h^{-1}$. The experiments were independently repeated three times, and the data for one time are shown here.

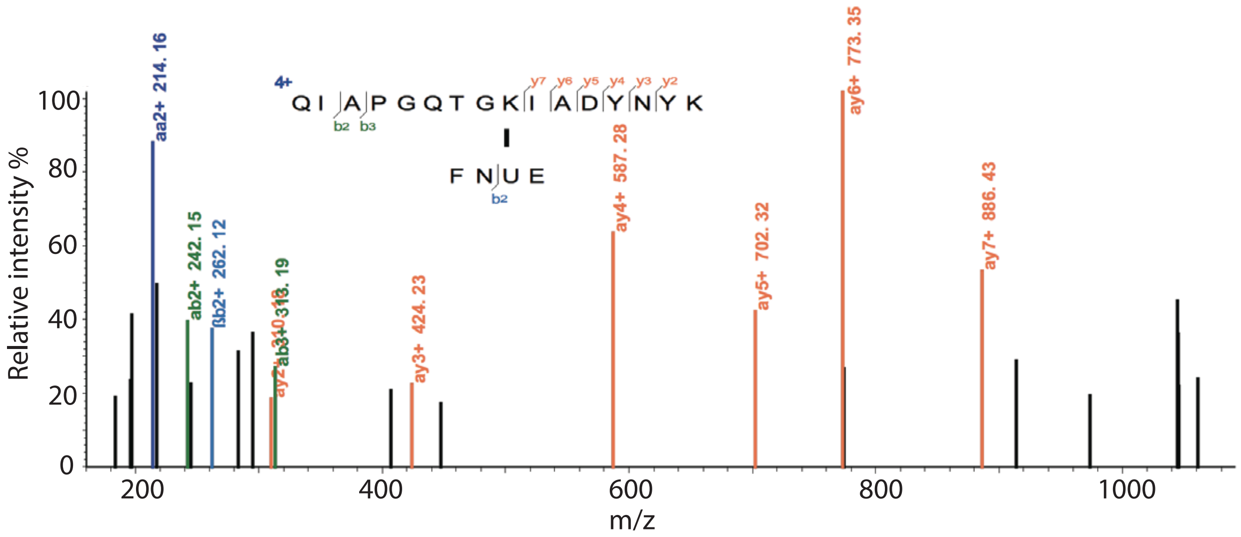


**Figure S7.** Tandem mass spectrum of human ACE2(34FSY) cross-linking with the Spike RBD, confirming that FSY34 in ACE2(34FSY) cross-linked with Lys 417 in the Spike RBD. U represents FSY.
